# Optic nerve regeneration requires the intracellular domain of LIFRa/CD118

**DOI:** 10.1101/2025.10.14.682231

**Authors:** Qian Jiang, Cong Wang, Yuerong Ren, Peiyun Duan, Ke tian, Xiangwei Duan, Binghan Cai, Changzhong Xu, Jian Li, Larry Benowitz, Ningli Wang, Bing Jiang, Lili Xie

## Abstract

Identifying cell-autonomous and non-autonomous factors that govern retinal ganglion cells’ (RGCs) ability to extend axons is an important step in developing therapies to achieve recovery after optic nerve injury.

Here we report that the intracellular domain of the leukemia inhibitory factor receptor (LIFR/CD118) is essential for mature RGCs’ ability to regenerate injured axons independent of the cognate ligand (LIF) and other therapies. Overexpression of LIFR in adult RGCs induces neurite outgrowth in cultured RGCs and axon regeneration in vivo while strongly amplifying RGCs’ response to LIF itself and to unrelated growth factors. Conversely, in loss-of-function studies, down-regulation of LIFR eliminates the pro-regenerative effects of Pten deletion and other potent stimuli. LIFR bidirectionally regulates the constitutive activity of the MAP kinase pathway, in contrast to LIF itself, which primarily activates pSTAT3. Expression of a truncated LIFR lacking the extracellular domain is sufficient to promote axon regeneration and selectively increases the survival of particular RGC subtypes, placing the LIFR intracellular domain as an essential, cell-autonomous regulator of optic nerve regeneration.

## Introduction

In mammals, the regenerative ability and trophic responsiveness of retinal ganglion cells (RGCs) decline after birth(1–4), contributing the irreversible visual losses commonly seen after traumatic or ischemic optic nerve injury and in several neuro-ophthalmic conditions, including glaucoma. Along with other important changes known to occur over the course of RGC development(5–7), the downregulation of critical receptors for trophic factors could potentially play a role in limiting recovery. Because several previously unrecognized trophic factors and their cognate receptors have been shown to stimulate regeneration of the injured optic nerve and to synergize with other treatments(7–17), we have extended the search for other important ligands and receptors. Receptors are highly specialized biological transducers that receive signals and respond to outside stimuli, commonly the cognate ligands. By screening the multiple receptors found to be expressed in adult RGCs in our earlier RNA-seq analysis(7), we discovered that expression of the leukemia inhibitory factor receptor (LIFR) correlates with the decline of RGCs’ regenerative potential. LIFR belongs to the type I cytokine receptor family, acting as a receptor for LIF and the related proteins oncostatin M (OSM) and cardiotrophin 1 (CTF1)(18, 19). Although varying effects of LIF itself on optic nerve regeneration have been reported(9, 20, 21), the present study shows that the receptor, LIFR, is itself a key cell-autonomous regulator of optic nerve regeneration independent of the presence or absence of LIF, and is required for the activity of unrelated pro-regenerative treatments, bidirectionally gating the activity of essential signal transduction pathways and expression of multiple pro-and anti-regenerative genes in mature RGCs.

## Results

### 1. LIFR expression correlates with RGC growth potential

#### 1.1. LIFR expression after optic nerve injury and during regeneration

To investigate whether the expression of any receptors correlates with RGCs’ growth state, we first examined receptor expression profiles in RGCs after injury and under regenerative conditions. RGCs were FACS-sorted from mature mice at 1, 3 or 5 days post crush (dpc) with or without intraocular injections of zymosan, a yeast cell wall preparation that induces optic nerve regeneration by elevating intraocular inflammation (Figure S1D) and levels of immune cell-derived growth factors (*e.g.*, oncomodulin, SDF-1)(10–12, 22, 23). These studies used a mouse line in which RGCs were genetically labeled (vglut2-Cre+/-crossed with *ZsGreen^flx/flx^-STOP^flx/flx^-tdTomato*), as verified by immunostaining against the RGC marker βIII-tubulin (Figure S1A). Using quantitative polymerase chain reaction (qPCR) to assess expression profiles of 53 receptors previously found by RNAseq to be expressed in normal adult RGCs(7), only the LIF receptor (Lifr) showed a significant decrease after optic nerve crush (NC) and partial recovery with the pro-regenerative treatment (Table 1). In a follow-up study, we found that Lifr mRNA in mature RGCs was about half that seen at postnatal 5d (P5) (*p<*0.01, unpaired t-test, Figure S1B) and, in conformity with the results reported above, was further reduced by 57% from normal adult levels at 3 dpc (*p<*0.01) and by 97% at 5 dpc (*p<*0.0001, all one-way ANOVA) (Figure S1C). NC-induced reduction of LIFR mRNA was partially alleviated by intraocular zymosan (compared to NC-only groups, intravitreal zymosan enhanced LIFR mRNA levels in adult RGCs 3.0-fold at day 3, *p<*0.01; and 2.6-fold at day 5, *p<*0.01; all unpaired t-tests) (Figure S1D). At the protein level, compared to RGCs from mice at P1, LIFR intensity declined by ∼ 30% 7 days after birth (*p<*0.0001, one-way ANOVA: Figure 1A, B) but did not decrease further at later time points (all *p>*0.05, one-way ANOVA, Figure 1A, B, Figure S1E, F; RGCs immunostained for the RNA-binding protein with multiple splicing, RBPMS). In parallel with the decline in LIFR mRNA levels, optic nerve injury further reduced in mature RGCs by 25-32% (25.3% reduction at 1 dpc, *p<*0.05; 31.6% reduction at 5 dpc, *p<*0.01, all one-way ANOVA) (Figure 1A, B). The consistent disparity between the mRNA and protein changes suggests that the LIFR protein may turn over slowly.

**Figure 1.**
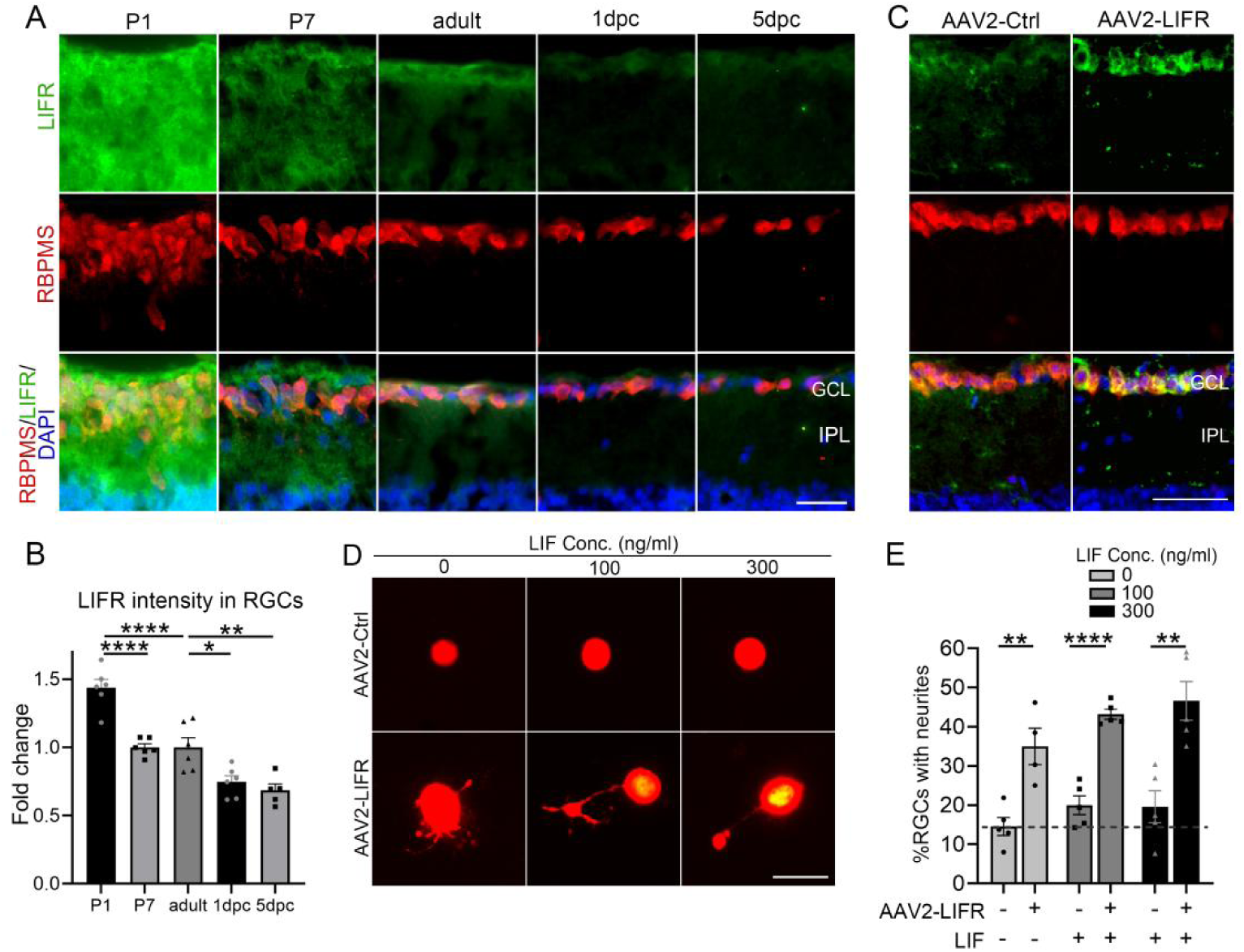
LIFR expression in RGCs. (A) Retinal cross-sections from P1, P7, and mature (6w) mice; and 1, 3, and 5 days post-crush (dpc) immunostained for LIFR (*green*), RBPMS (RGC pan-marker, *red*) and DAPI (*blue*). GCL: ganglion cell layer, IPL: inner plexiform layer. Scale bar = 30 µm. (B) Quantification of LIFR intensity in RGCs from mice at time points shown in (A) (\**p<*0.05, \*\**p<*0.01, \*\*\*\**p<*0.0001; n = 5-6 eyes per group). (C) Retinal cross-sections from mice two weeks following AAV2-Ctrl or AAV2-LIFR injection immunostained with antibodies to LIFR (*green*) and RBPMS (*red*) and DAPI (*blue*). Scale bar = 30 µm. (D) Representative images showing neurite outgrowth in cultured RGCs (*red*) from vglut2-tdTomato mice with indicated treatments at day 3. Scale bar = 50μm. (E) Quantitation of neurite outgrowth in RGCs (\*\**p<*0.01, \*\*\*\**p<*0.0001; n = 4-5 wells per group). Bars show means ± SEM.

**Table 1.** QPCR screening for 53 receptors in adult RGCs.

| Genes | NC(1d) | NC+zymo(1d) | NC(3d) | NC+zymo(3d) | NC(5d) | NC+zymo(5d) |
| --- | --- | --- | --- | --- | --- | --- |
| *Lifr* | *n.s.* | *n.s.* | -(**) | +(**) | -(***) | +(**) |
| *Gfra4* | +(***) | *n.s.* | *n.s.* | *n.s.* | *n.s.* | *n.s.* |
| *Amfr* | +(***) | NA | *n.s.* | *n.s.* | *n.s.* | *n.s.* |
| *Ifngr2* | NA | *n.s.* | +(*) | *n.s.* | *n.s.* | *n.s.* |
| *Crll* | NA | *n.s.* | *n.s.* | *n.s.* | +(*) | *n.s.* |
| *Cxcr4* | *n.s.* | *n.s.* | *n.s.* | *n.s.* | *n.s.* | -(*) |

Of 53 receptors found to be expressed in RGCs, only 6 showed statistically significant changes following axonal injury with or without a pro-regenerative treatment (intraocular zymosan). NA, *n.s.*, \*, \*\*, and \*\*\* indicate undetermined, $p > 0.05$ , $p < 0.05$ , $p < 0.01$ , and $p < 0.001$ respectively. + and – indicate upregulated and downregulated respectively.

To characterize developmental changes in LIFR expression in greater detail, we assessed LIFR protein levels in rats at embryonic day 17 (E17), E21, P1, P2, P8 and P11, time points previously used to study developmental changes in RGCs’ growth potential(24). In conformity with the finding that embryonic RGCs have a stronger intrinsic capacity for axon growth than postnatal RGCs(24), we saw a 16.5% decrease in LIFR expression between E17 and E21 that remained at a similar level through P2 (at P1 and P2, *p*< 0.05, one-way ANOVA), then declined further by P8 (55.9% decline relative to E17 at P8, 52.2% decline at P11, both *p*<0.0001, one-way ANOVA: Figure S1G, H).

#### 1.2. LIFR overexpression is sufficient to stimulate neurite outgrowth in adult RGCs

To evaluate the efficacy of LIFR overexpression, we expressed the *Lifr* gene via the RGC-selective adeno-associated virus serotype 2.2 (AAV2-*Lifr*; may also infect some amacrine cells and glia(25–27)) which enhanced LIFR protein levels 2.2-fold in RBPMS+ RGCs (compared to the control group: *p<*0.01, unpaired t-test, Figure 1C and Figure S1I). Next, to test the consequences of LIFR overexpression, we again overexpressed *Lifr* (*vs.* a control construct) in adult vGLUT2-tdTomato mice (in which RGCs are genetically labeled with tdTomato), then prepared dissociated retinal cultures two weeks later. After 3 days in culture, mature RGCs overexpressing LIFR showed a 1.4-fold increase in neurite outgrowth (*p<*0.01, unpaired t-tests: Figure 1D, E). Regarding the cognate ligand, in the absence of receptor overexpression, LIF at either 100 or 300 ng/ml had no detectable effect on neurite outgrowth nor RGC survival (*p>*0.05, Figure 1D, E, Figure S2). However, with LIFR overexpression, LIF approximately doubled levels of neurite outgrowth: 1.2-fold increase with LIF at 100 ng/ml (*p<*0.0001) and 1.4-fold increase with LIF at 300 ng/ml (*p<*0.01; all unpaired t-tests; Figure 1D, E). In one case, we detected a 24.3% decrease in RGC survival in the presence of the ligand (AAV2-*Lifr* + LIF, 100 ng/ml, *p<*0.05, unpaired t-test, Figure S2).

#### 1.3. Neutrophils express abundant LIF protein in response to intraocular zymosan

Because neutrophils recruited by intraocular zymosan secrete trophic factors that promote optic nerve regeneration(11, 12, 22), we tested whether these cells express LIF. By immunochemistry for the neutrophil marker Gr1, we confirmed the presence of numerous neutrophils in the posterior chamber of the eye from 12h to 3 d after intraocular Zymosan which decline by 5d (Figure S3A). We isolated neutrophils from the vitreous using magnetic beads coated with an anti-Ly6G antibody at different time points following intraocular zymosan and assessed neutrophil purity by flow cytometry after incubation with an anti-Gr1 primary antibody conjugated with FITC. Neutrophils, characterized by Giemsa-stained multilobed nuclei, accounted for 93.8% of the isolated cells (Figure S3B). Because the posterior chamber of the eye normally contains very few cells, we instead quantified absolute mRNA levels. Compared to the sham-treated group, LIF mRNA levels increased dramatically within the first 5 days after zymosan injection (49.4-fold increase at day 1, *p>*0.05; 213.8-fold increase at 3d, *p<*0.0001; 51.6-fold increase at 5d, *p>*0.05, all one-way ANOVA: Figure S3C). Thus, infiltrative neutrophils express LIF in addition to the trophic factors reported previously(8, 9, 11, 23).

## 2. LIFR overexpression induces optic nerve regeneration independent of its cognate ligand

### 2.1. AAV2-LIFR promotes axon regeneration

To investigate the effect of LIFR overexpression *in vivo*, we injected AAV2-LIFR intraocularly into adult mice two weeks prior to NC. Even in the absence of exogenous LIF, LIFR overexpression augmented axon regeneration 3.5-fold over mice receiving a control vector when evaluated 2 weeks after nerve injury (*p<*0.0001, unpaired t-test, Figure 2A, B) without altering RGC survival (*p>*0.05, unpaired t-test, Figure 2A, C). As previously reported(7, 10, 11, 28–30), RGC survival in whole-mounted retina was visualized by immunostaining against β III-tubulin (antibody TUJ1). We verified that TUJ1 staining was only seen in RBPMS-positive RGCs (Figure S4A).

**Figure 2.**
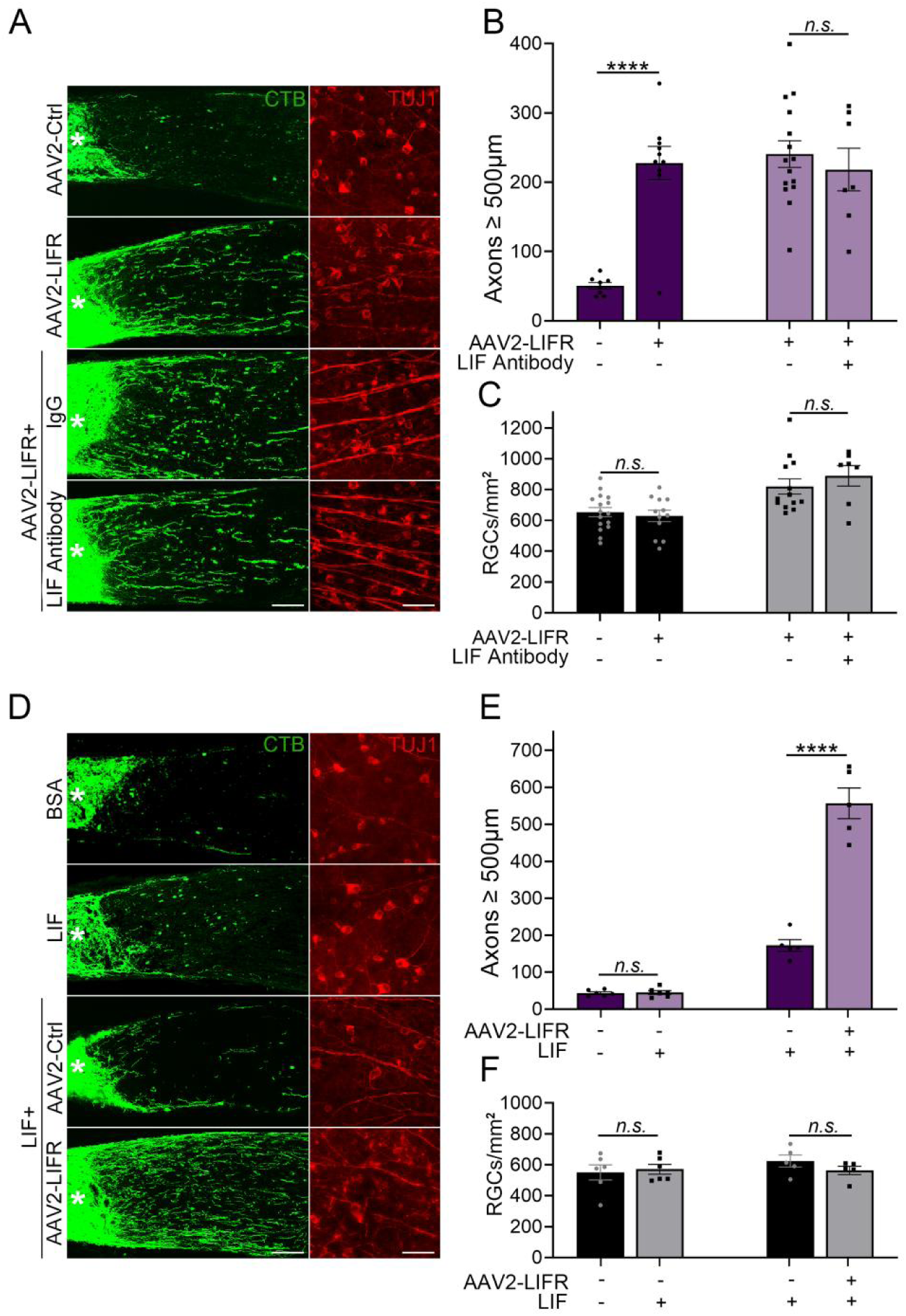
LIFR overexpression promotes axon regeneration in the mature optic nerve. (A) Regenerating axons and RGC survival in whole-mounted retinas visualized by fluorescent CTB (*green*) and immunostaining against βIII-tubulin (*red*), respectively, two weeks after NC. Mice received intraocular injections of AAV2-Ctrl, AAV2-LIFR, AAV2-LIFR combined with an isotype control antibody, or AAV2-LIFR combined with LIF neutralizing antibody. An asterisk indicates the injury site. Scale bar = 100 µm in the left panel, 50 µm in the right panel. (B) Quantification of regenerating axons 500 µm from the injury site (*n.s.*: *p>*0.05, \*\*\*\**p<*0.0001; n = 7-15 nerves per group). (C) Quantification of RGC survival (*n.s.: p>*0.05; n = 7-16 eyes per group). (D) Representative images for regenerating axons and RGC survival in whole-mounted retinas with indicated treatment. The asterisk indicates the injury site. Scale bar = 100 µm in the left panel, 50 µm in the right panel. (E) Quantification of regenerating axons (*n.s.*: *p>*0.05, \*\*\*\**p<*0.0001; n = 5-6 nerves per group). (F) Quantification of RGC survival (*n.s.*: *p>*0.05; n = 5-6 eyes per group). Bars show means ± SEM.

### 2.2. The effect of LIFR overexpression is ligand-independent

To determine whether the effect of LIFR overexpression requires the presence of LIF, the cognate ligand, we introduced a LIF-neutralizing antibody to diminish endogenous levels of the protein. LIF concentration in aqueous humor and serum is ∼4 pg/ml (0.15 pM: human)(31), and we used a >10,000-fold excess of LIF neutralizing antibody (400 ng/ml, ∼2.7 nM) to reduce levels of the protein. As before, AAV2-*Lifr* was introduced intraocularly two weeks before NC, and the LIF neutralizing antibody (400 ng/ml) was injected intravitreally twice, once at 5d before and again immediately after NC. The LIF neutralizing antibody did not alter AAV2-*Lifr*-induced axon regeneration (*p>*0.05, unpaired t-test, Figure 2A, B) and had no impact on RGC survival (*p>*0.05, unpaired t-test, Figure 2A, C). We also tested the effect of applying neutralizing antibodies against two alternative ligands for LIFR, cardiotrophin-1 (anti-CTF1; 400 ng/ml) and oncostatin M (anti-OSM; 3000 ng/ml) in the vitreous. Neither antibody altered AAV2-LIFR–induced axon regeneration (*p*>0.05, one-way ANOVA, Figure S4B-D).

### 2.3. LIFR overexpression augments the effects of LIF

In complementary studies, we investigated whether LIFR overexpression improves RGCs’ response to the cognate ligand. Whereas a single intraocular injection of LIF (1 µg/eye) immediately after NC had no detectable effect (*p>*0.05 compared to BSA, 1 µg/eye, unpaired t-test, Figure 2D, E), it had a clear effect when LIFR was overexpressed (intraocular AAV2-*Lifr*: 2.2-fold increase compared to LIF plus AAV2-Ctrl, Figure 2D, E; 1.4-fold increase compared to AAV2-*Lifr*: both *p*< 0.0001, unpaired t-tests, Figure 2A, B, D, E). In all cases, RGC survival was unaffected (*p>*0.05, unpaired t-test, Figure 2A, C, D, F).

## 3. AAV2-LIFR enhances RGCs’ response to unrelated growth factors

We next examined whether receptor overexpression also increases RGCs’ responsiveness to unrelated trophic factors, including insulin-like growth factor 1 (IGF1, 1 µg/eye), stromal cell-derived factor 1 (SDF1, 0.9 µg/eye) or oncomodulin (Ocm, 90 ng/eye), all of which have been shown to promote optic nerve regeneration(8, 9, 11, 13). As reported earlier(8, 9, 11), single intraocular injections of either IGF1 or SDF1 stimulated moderate levels of axon regeneration (IGF-1: *p<*0.01, Figure 3A, B; SDF-1, *p<*0.0001, unpaired t-tests, Figure 3A, D), and strongly increased RGC survival (72.4% increase for IGF-1, *p<*0.001, Figure 3C; 81.3% increase for SDF-1, *p<*0.0001, unpaired t-tests, Figure 3E), whereas Ocm (plus CPT-cAMP) had little effect in the absence of complementary factors (*e.g*. Pten deletion, SDF1 expression)(9) (*p>*0.05, unpaired t-test, Figure 3F). However, when combined with LIFR overexpression, all three factors roughly doubled levels of axon regeneration compared to receptor overexpression alone (*p<*0.05, one-way ANOVA, Figure 3A, B, D, F). Thus, LIFR elevation strongly enhances RGCs’ responsiveness to both the cognate ligand and to unrelated trophic factors. The combinations did not improve RGC survival above the levels achieved with individual factors alone (*p>*0.05, one-way ANOVA, Figure 3A, C, E, G).

**Figure 3.**
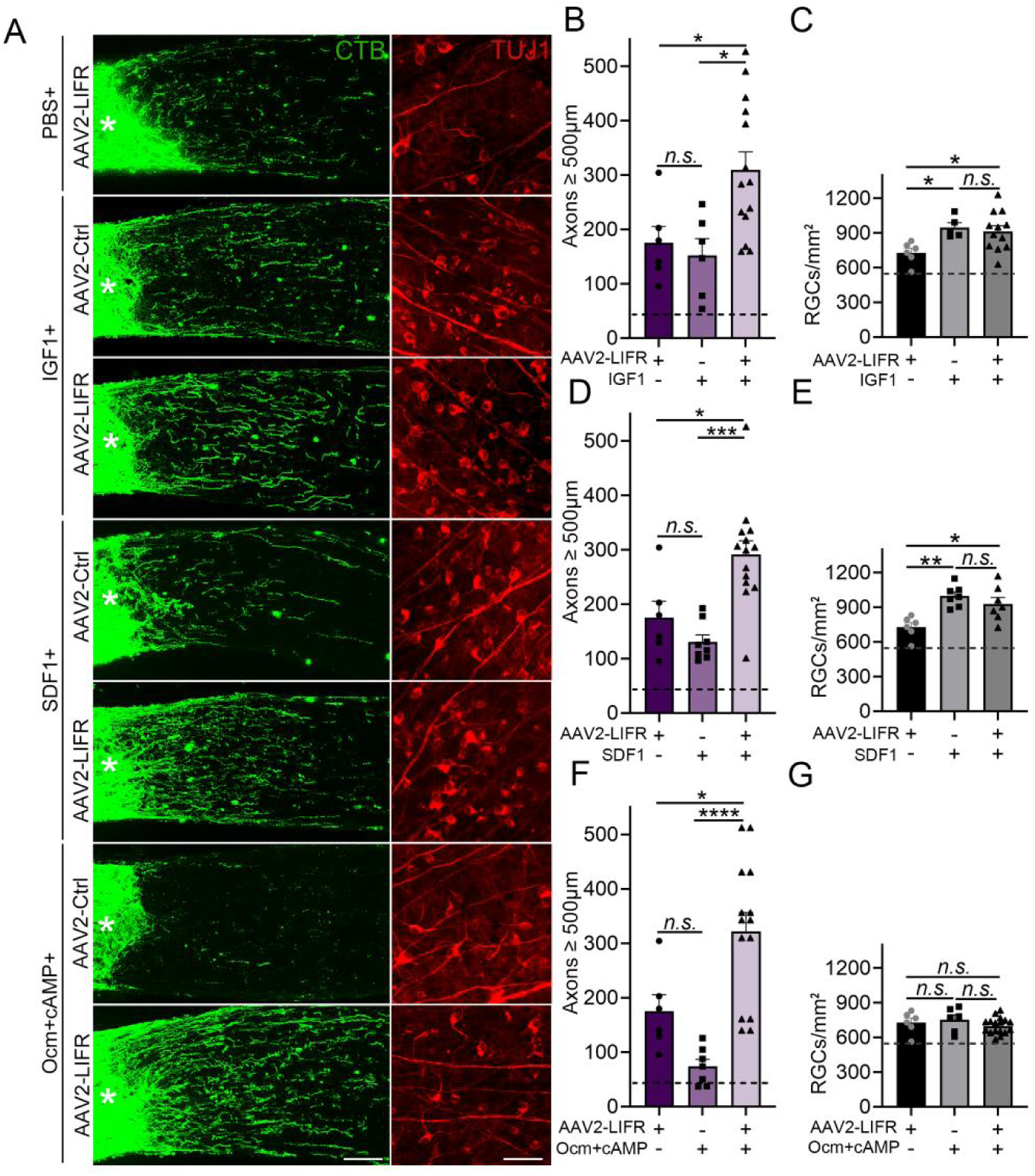
AAV2-LIFR enhances RGCs’ responsiveness to unrelated trophic factors. (A) Regenerating axons and RGC survival visualized by fluorescent CTB transport (*green*) and immunostaining against βIII-tubulin (*red*), respectively, two weeks after NC with indicated treatments. Asterisk indicates injury site. Scale bar = 100 µm in the left panel, 50 µm in the right. (B) Quantification of regenerating axons (AAV2-*Lifr* +IGF1 *vs.* AAV2-Ctrl+IGF1). Dashed line indicates the level of axon regeneration in the BSA-treated control group. (*n.s.*: *p>*0.05, \**p<*0.05, n = 6-14 nerves per group). (C) Quantification of RGC survival. Dashed line indicates RGC survival in the BSA-treated group. (*n.s.*: *p>*0.05, \**p<*0.05; n = 5-12 eyes per group). (D) Quantification of axon regeneration. (*n.s.*: *p>*0.05, \**p<*0.05, \*\*\**p<*0.001; n = 6-14 nerves per group). (E) Quantification of RGC survival. (*n.s.*: *p>*0.05, \**p<*0.05, \*\**p<*0.01; n = 6-7 eyes per group). (F) Quantification of axon regeneration. Dashed line indicates baseline regeneration in BSA-treated group. (*n.s.*: *p>*0.05, \**p<*0.05, \*\*\*\**p<*0.0001; n = 6-14 nerves per group). (G) Quantification of RGC survival. Dashed line indicates RGC survival in the BSA-treated group (*n.s.*: *p>*0.05; n = 6-17 eyes per group). Bars show mean ± SEM.

## 4. Loss of function: Optic nerve regeneration requires LIFR

To investigate the necessity of LIFR for optic nerve regeneration, the receptor was knocked down in RGCs of mature mice using an AAV2 expressing a targeted small hairpin RNA (shRNA). After two weeks, immunohistochemistry confirmed a reduction in LIFR intensity in RBPMS+ RGCs to background, *i.e*., the level seen when the primary antibody was omitted during immunostaining (*p<*0.05, unpaired t-test, Figure 4A, B). AAV2-shLIFR reduced zymosan-induced axon regeneration to the baseline level seen after NC alone (*p<*0.0001, unpaired t-test, Figure 4C, D), and somewhat unexpectedly, deceased zymosan-induced RGC survival to baseline level (*p<*0.001, unpaired t-test, Figure 4C, E). To explore the importance of LIFR more generally, we also tested the effect of AAV2-shLIFR on regeneration induced by deletion of phosphatase and tension homolog (Pten), a negative regulator of the PI3 kinase/Akt pathway that is perhaps the most potent single pro-regenerative treatment reported to date(32–34). AAV2-shLIFR fully abolished both axon regeneration and the neuroprotective effect of Pten deletion (*p<*0.001, unpaired t-test, Figure 4F-H).

**Figure 4.**
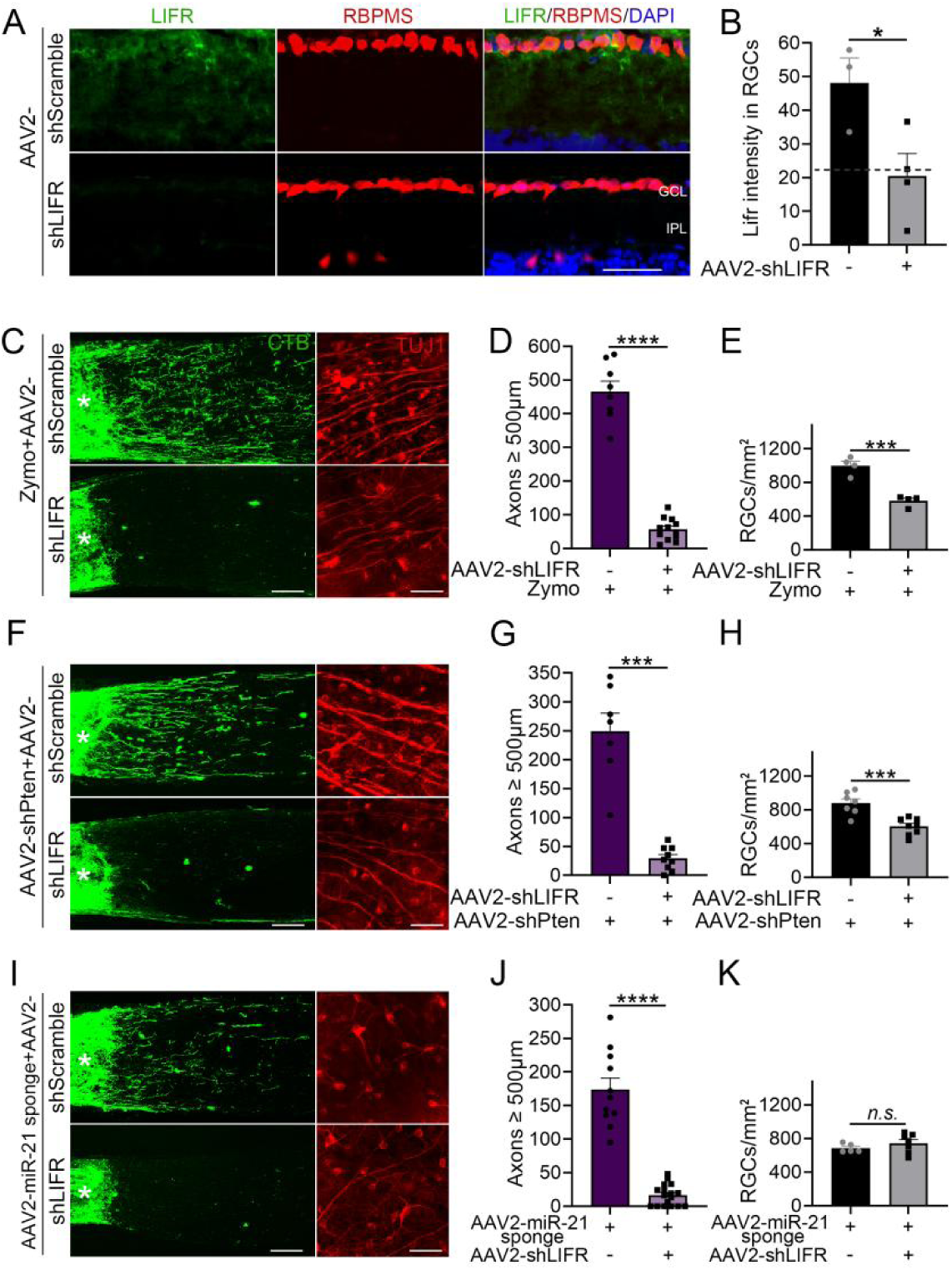
LIFR deletion abolishes optic nerve regeneration. (A) Retinal cross-sections from mice two weeks after AAV2-sh-Scramble or AAV2-shLIFR injection immunostained for LIFR (*green*) and RBPMS (*red*); and stained with DAPI (*blue*). GCL: ganglion cell layer, IPL: inner plexiform layer. Scale bar = 30 µm. (B) Quantification of LIFR intensity in RGCs. Dashed line indicates LIFR intensity when the 1st antibody is omitted during immunostaining (\**p<*0.05; n = 3-4 eyes per group). (C) Representative images showing axon regeneration (*green*) and RGC survival (*red*). Asterisk indicates the injury site. Scale bar = 100 µm in the left panel, 50 µm in the right panel. (D) Quantification of axon regeneration (\*\*\*\**p<*0.0001; n = 8-12 nerves per group). (E) Quantification of RGC survival (AAV2-shLIFR+zymo *vs.* AAV2-shScramble+zymo, \*\*\**p<*0.001; n = 4 eyes per group). (F) Fluorescent CTB (*green*) labeled regenerating axons in nerve sections and βIII-tubulin (*red*) labeled RGC survival in whole-mounted retina with indicated treatments. Asterisk indicates the injury site. Scale bar = 100 µm in the left panel, 50 µm in the right panel. (G) Quantification of axon regeneration. (AAV2-shLIFR + AAV2-shPten *vs.* AAV2-shScramble + AAV2-shPten, \*\*\**p<*0.001; n = 7-9 nerves per group). (H) Quantification of RGC survival (AAV2-shLIFR+AAV2-shPten *vs.* AAV2-shScramble+AAV2-shPten, \*\*\**p<*0.001; n = 7-8 eyes per group). (I) Regenerating axons and RGC survival visualized by fluorescent CTB (*green*) and immunostaining against βIII-tubulin (*red*), respectively, two weeks after NC with indicated treatments. Scale bar = 100 µm in the left panel, 50 µm in the right panel. (J) Quantification of axon regeneration (AAV2-shLIFR + AAV2-miR-21 sponge *vs.* AAV2-shScramble + AAV2-miR-21 sponge, \*\*\*\**p<*0.0001; n = 11-17 nerves per group). (K) Quantification of RGC survival (AAV2-shLIFR + AAV2-miR-21 sponge *vs.* AAV2-shScramble + AAV2-miR-21 sponge, *n.s.*: *p>*0.05; n = 5-7 eyes per group). Bars show means ± SEM.

MicroRNAs (miRNAs) are small noncoding RNAs that play important regulatory roles by targeting mRNAs for cleavage or translational repression(35, 36). Mir-21 has been reported to interact with LIFR mRNA directly(37, 38), which we verified by showing that mir-21 binds to the 3’ untranslated region (UTR) of LIFR mRNA (dual-Luciferase reporter assay: Figure S5A, B). We found that miR-21 is highly expressed in RBPMS+ RGCs using fluorescence *in situ* hybridization (FISH) (Figure S5C). We next constructed an AAV2 expressing miR-21 antisense sequences (AAV2-miR-21-sponge) to sequester miR-21 away from its endogenous targets. Like LIFR overexpression, AAV2-mir-21-sponge enhanced axon regeneration 3.5-fold over baseline (*p<*0.0001 compared to AAV2-Ctrl-sponge, unpaired t-test, Figure S5D, E) with no impact on RGC survival (*p>*0.05, unpaired t-test, Figure S5D, F). However, when LIFR expression was knocked down (intraocular AAV2-shLIFR 3 weeks before optic nerve crush) the effect of AAV2-miR-21-sponge (intraocular, 2 weeks before optic nerve crush), was abolished (*p<*0.0001, unpaired t-test, Figure 4I, J), confirming the necessity of LIFR for RGC axon regeneration.

## 5. Transcriptomic analyses

To investigate the molecular consequences of LIFR over-expression, RGCs from vglut2-tdTomato mice were FACS-isolated two weeks after intraocular injection of AAV2-*Lifr* or AAV2-Ctrl, then analyzed by bulk RNA-seq. AAV2-*Lifr* had pronounced transcriptional effects on adult RGCs, as indicated by principal component analysis (PCA, Figure S6A) and the vast number of differentially expressed genes (DEGs) (4020 significant gene changes, q<0.05, log2|FC|>1, 2112 up-regulated and 1908 down-regulated, AAV2-LIFR *vs.* AAV2-Ctrl, Figure 5A, Figure S6C). AAV2-*Lifr* upregulated genes encoding many growth factors (*e.g*., Egf, Ngf, Igf2, Ccl5) and receptors, (Igf1r, Igf2r) along with the expected increase in LIFR, while downregulating the expression of genes associated with inhibition of regeneration (Rtn4r (Nogo receptor, NgR), Rtn4rl2 (NgR2), Lingo1, Cspg5), growth cone collapse, (Sema6c, Sema4f) and cell death (Nlrp1a, Faf1) (Figure 5B). Gene Ontology (GO) enrichment analysis revealed that LIFR overexpression upregulated groups of genes relevant to positive regulation of the MAPK cascade, positive responses to cytokine stimulus, cytokine receptor activity, positive regulation of axon regeneration, and regulation of receptor signaling pathways via JAK-STAT and downregulation of groups of genes associated with autophagic cell death, programmed necrotic cell death, intrinsic apoptotic signaling, and negative regulation of axonogenesis, axon extension, and regeneration (Figure 5C).

**Figure 5.**
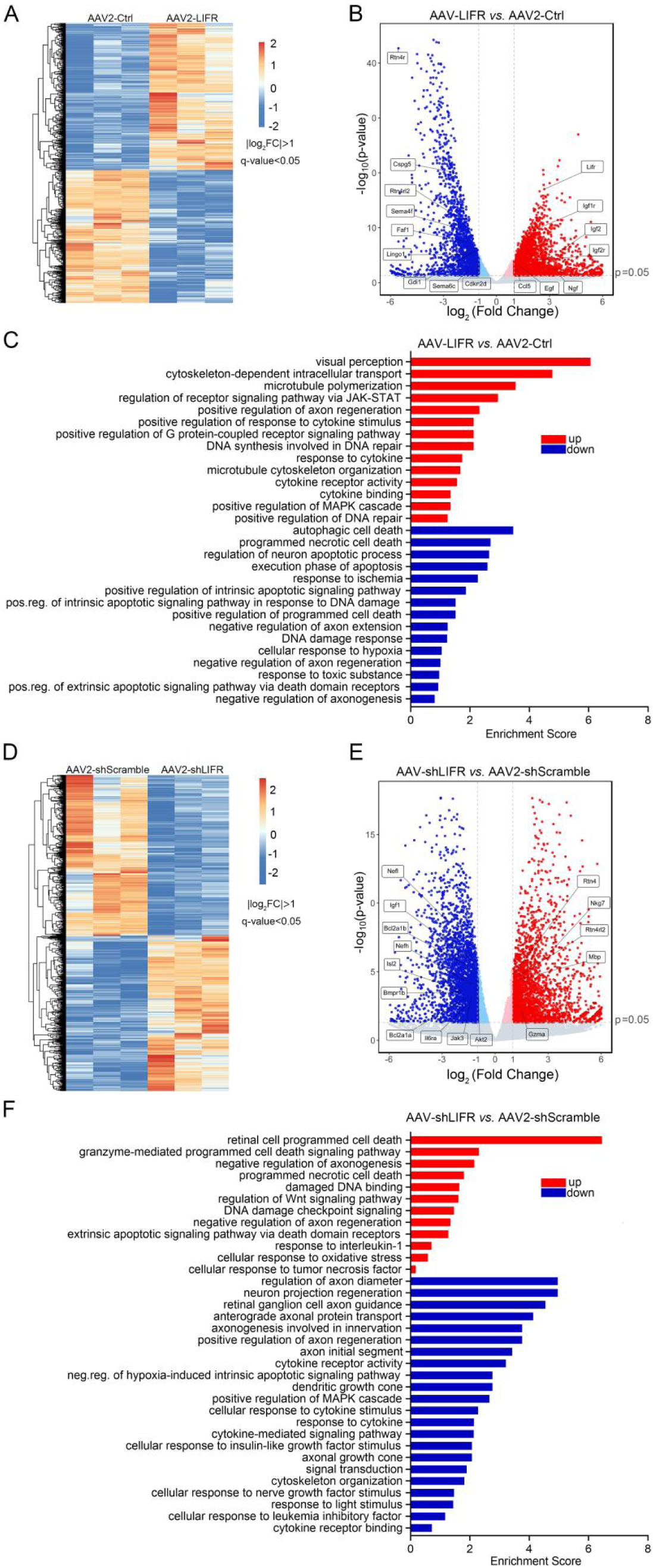
Transcriptional effects. (A, D) Heatmap showing the magnitude of up-regulated (*orange to red*) and down-regulated (*blue*) genes (q<0.05, log_2_|FC|>1) in RGCs two weeks following intraocular AAV2-LIFR (A, AAV2-LIFR *vs.* AAV2-Ctrl) or AAV2-shLIFR (D, AAV2-shLIFR *vs.* AAV2-shScramble, 3 replicates in each group). (B, E) Volcano plot showing individual up-regulated (*red*) and down-regulated (*blue*) genes in RGCs following transfection with AAV2-LIFR (B, AAV2-LIFR *vs.* AAV2-Ctrl) or AAV2-shLIFR (E, AAV2-shLIFR *vs.* AAV2-shScramble). (C) GO enrichment analysis of differentially expressed genes with indicated comparison in RGCs. Up-regulated (*red*) and down-regulated (*blue*) GO terms (AAV2-LIFR *vs.* AAV2-Ctrl). (F) GO enrichment analysis of differentially expressed genes with indicated comparison in RGCs. Up-regulated (*red*) and down-regulated (*blue*) GO terms following AAV2-shLIFR transfection (n = 3 replicates per group, AAV2-shLIFR *vs.* AAV2-shScramble).

In complementary studies, we explored the transcriptional effect of knocking down LIFR (via AAV2-shLIFR) in RGCs, again using bulk RNA-seq. As with LIFR overexpression, LIFR knock-down had robust transcriptional effects as revealed by PCA, heatmap analysis (Figure S6B, Figure. 5D), and the large number of DEGs (4708 genes, q<0.05, log2|FC|>1, 2065 up-regulated and 2643 down-regulated, AAV2-shLIFR *vs.* AAV2-shScramble, Figure 5D, Figure S6D). In a reversal of the effects of LIFR overexpression, LIFR knock-down enhanced the expression of myelin-associated inhibitors of axon growth and their receptors (Rtn4 (Nogo), Rtn4r (NgR) and Mbp) while downregulating anti-apoptotic genes (Bcl2a1a, Bcl2a1b, Bmpr1b) and genes encoding growth factors, receptors, and their effectors (Igf1, Isl2, Il6ra, Nefl, Nefh, Akt2, and Jak3) (Figure 5E). At the same time, GO enrichment analysis showed that LIFR knock-down led to the upregulation of groups associated with programmed cell death, damaged DNA binding, and negative regulation of axonogenesis and axon regeneration, and downregulation of terms relevant to the positive regulation of MAPK cascade, cellular response to Igf and Ngf (and LIF), cytokine-mediated signaling pathways, positive regulation of axon regeneration, and RGC axon guidance (Figure 5F).

## 6. Supporting evidence for LIFR regulation of signal transduction pathways

We further investigated the role of LIFR in neuroprotective and pro-regenerative signal pathways by immunohistochemistry. As before, we manipulated levels of LIFR expression by intraocular injections of either AAV2-*Lifr* or AAV2-shLIFR, then evaluated levels of phosphorylated (active) extracellular signal-regulated kinase (pERK), phosphorylated protein kinase B (pAKT), phosphorylated ribosomal protein S6 (pS6), and phosphorylated signal transducer and activator of transcription 3 (pSTAT3) in RBPMS+ RGCs two weeks later using appropriate antibodies. In conformity with the transcriptomic results, LIFR overexpression increased pErk levels in RGCs (*p<*0.01 compared to the AAV2-Ctrl group, unpaired t-test, Figure 6A and B), whereas LIFR knock-down reduced pErk levels in RGCs (by 33.7%: *p<*0.01 compared to AAV2-shScramble, unpaired t-test, Figure 6A and B). AAV2-shLIFR also reduced levels of pAKT and pS6, two markers of mTOR pathway activation, by 45.7% and 47.0% respectively (all *p<*0.001 compared to the AAV2-shScramble group, unpaired t-test, Figure 6A, C, D). LIFR overexpression also seemed to reduce pAKT and pS6 levels, though this did not attain statistical significance (*p>*0.05, unpaired t-test, Figure 6A, C, D). No significant differences were observed for pSTAT3 levels in RGCs as a result of LIFR overexpression or knockdown (*p>*0.05, unpaired t-test, Figure 6E, F).

**Figure 6.**
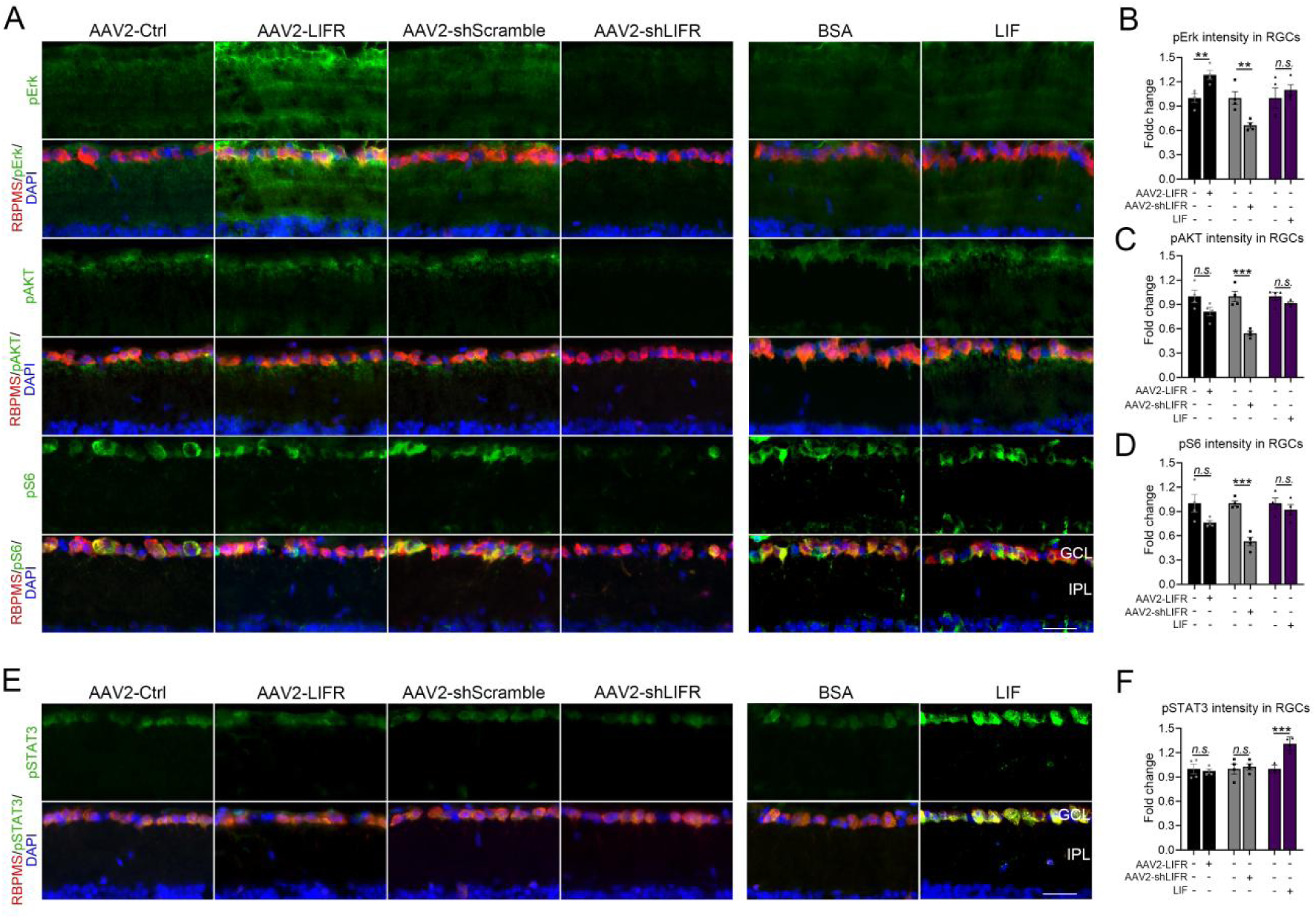
LIFR activates cell-signaling pathways related to CNS axon regeneration. (A) Retinal cross-sections from mice two weeks after intraocular AAV2-Ctrl, AAV2-*Lifr*, AAV2-shScramble, AAV2-shLIFR, BSA or LIF injection immunostained for pErk (*green*, up), pAKT (*green*, middle), pS6 (*green*, bottom), RBPMS (*red*), and DAPI (*blue*). GCL: ganglion cell layer, IPL: inner plexiform layer. Scale bar = 30 µm. (B) Quantification of pErk intensity in RBPMS+ RGCs with indicated treatments (\*\**p<*0.01; n=4 per group). (C) Quantification of pAKT intensity in RGCs with indicated treatments (*n.s.*: *p>*0.05, \*\*\**p<*0.001; n=4 per group). (D) Quantification of pS6 intensity in RGCs with indicated treatments (*n.s.*: *p>*0.05, \*\*\**p<*0.001; n=4 per group). (E) Retinal cross-sections from mice two weeks after intraocular AAV2-Ctrl, AAV2-Lifr, AAV2-shScramble or AAV2-shLIFR, BSA or LIF immunostained for pSTAT3 (*green*), RBPMS (*red*), and DAPI (*blue*). GCL: ganglion cell layer, IPL: inner plexiform layer. Scale bar = 30 µm. (F) Quantification of pSTAT3 intensity in RGCs with indicated treatments (*n.s.*: *p>*0.05, \*\*\**p<*0.001; n=4 per group). Bars show means ± SEM.

To compare the effects of LIFR overexpression with those of LIF itself, we also evaluated expression of the above signal molecules by immunohistochemistry 3 days after intraocular injections of recombinant LIF protein (1µg/eye). Intraocular LIF did not alter pERK, pAKT, or pS6 expression (compared to BSA control, *p*>0.05, unpaired t-test, Figure 6A-D), whereas, as expected, it increased pSTAT3 levels (*p*<0.001, unpaired t-test, Figure 6E, F). This finding suggests the possibility that LIFR overexpression and LIF might activate distinct downstream signaling pathways, although the mechanisms involved are unclear.

## 7. The intracellular domain of LIFR is sufficient to promote optic nerve regeneration

Because our experiments show that LIFR acts independently of its cognate ligands, we investigated whether a truncated form of the receptor lacking extracellular ligand binding sites is sufficient to stimulate regeneration. For this, we designed an AAV vector expressing a truncated LIFR (AAV2-LIFR Δ) that includes only the signal peptide, transmembrane and intracellular domains. Two weeks after intraocular injection of AAV2-LIFRΔ, we injured the optic nerve and evaluated regeneration after another two weeks. AAV2-LIFRΔ elevated axon regeneration 2.2-fold compared to AAV2-Ctrl (*p*<0.001, one-way ANOVA, Figure 7A, B), which was slightly but not significantly less than overexpression of the full-length receptor (26.0% lower compared to AAV2-LIFR: *p*>0.05, one-way ANOVA, Figure 7A, B). AAV2-LIFRΔ likewise did not affect RGC survival (*p*>0.05, one-way ANOVA, Figure 7A, C).

**Figure 7.**
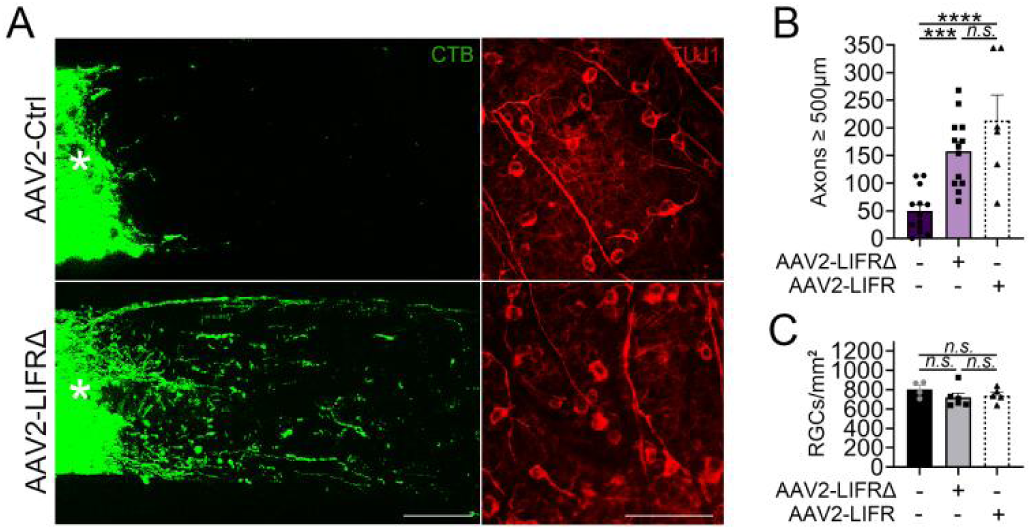
LIFR Intracellular Domain Promotes Regeneration (A) Representative images of regenerating axons (CTB, *green*) and RGCs survival (TUJ1, *red*) in mice treated with AAV2-LifrΔ or AAV2-Ctrl. Asterisks indicate the injury site. Scale bars: 100 µm for left panels, 75 µm for right panels. (B) Quantification of axon regeneration (*n.s.*: *p*>0.05, \*\*\**p*<0.001, \*\*\*\**p*<0.0001; n=6–13 nerves per group). (C) Quantification of RGC survival (*n.s.*: *p*>0.05; n=4–6 retinas per group) Bars show means ± SEM.

We also explored the role of LIFR on different RGC subtypes by immunostaining for SMI32 (against NF-H for aRGCs), CART (for on-off direction-selective RGCs, ooDSGCs), and melanopsin (for M1 - M5 intrinsically photosensitive RGCs, ipRGCs). Immunohistochemistry verified that all three markers colocalized with the pan-RGC marker, RBPMS (Figure S7A). Again, AAV2-LIFR Δ was intraocularly injected two weeks before nerve injury, and the survival of the three RGC subtypes was assessed after another 1 or 2 weeks. Despite the absence of an overall effect on RGC survival, AAV2-LIFRΔ strongly increased ipRGC survival (by 47.6% at 1 wk and 48.3% at 2 wk post-injury, respectively; both *p*<0.05, unpaired t-test, Figure S7B, C); decreased the survival of ooDSGC (by 40.4% at 1 wk and 41.9% at 2 wk post-injury; *p*<0.01, *p*<0.05, respectively, unpaired t-tests, Figure S7B, D) and did not affect the survival of αRGCs (*p*>0.05, unpaired t-test, Figure S7B, E). Thus, LIFR Δ appears to alter RGC survival in a subtype-specific way. Figure 8 schematically illustrates our major findings.

**Figure 8.**
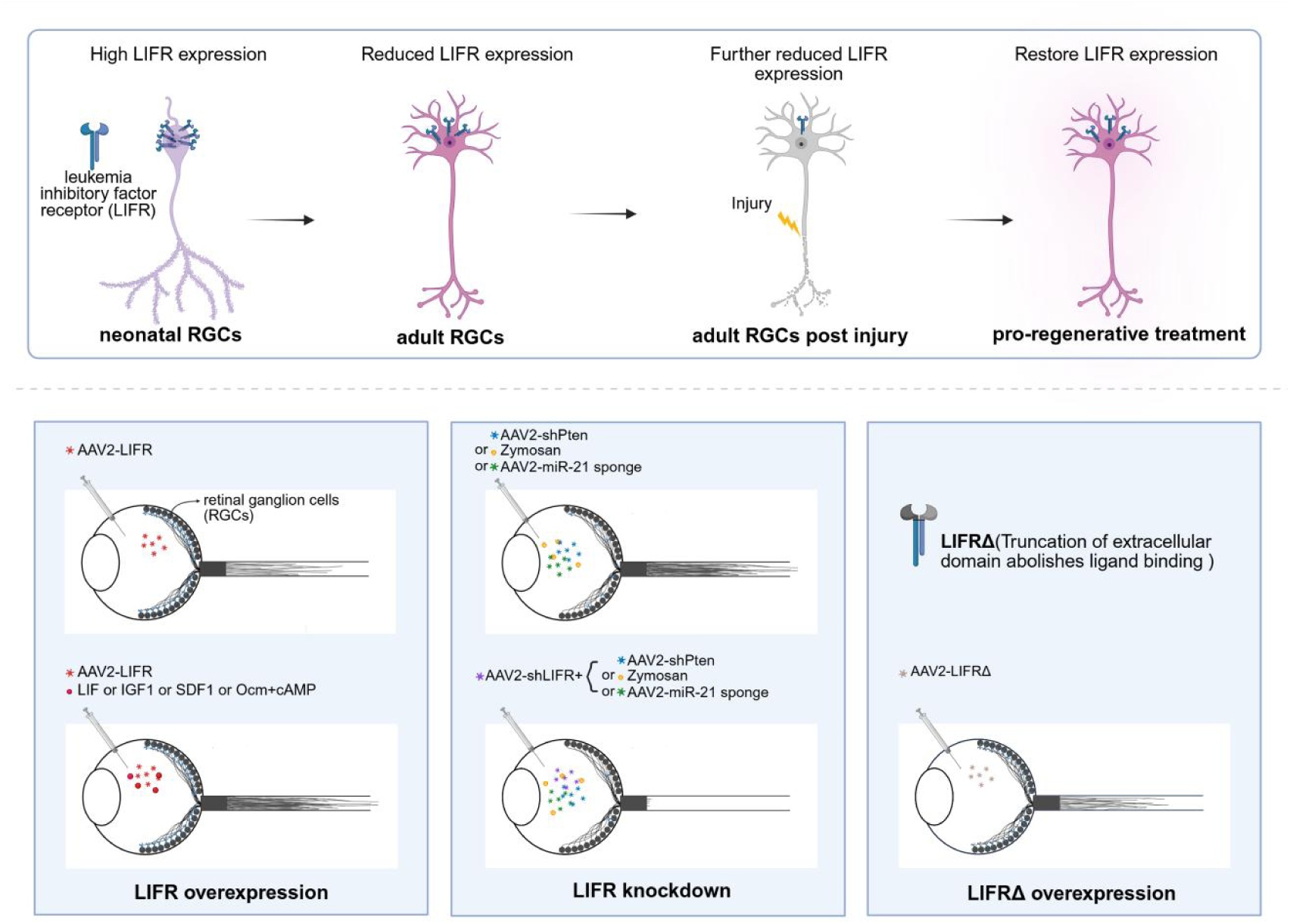
Schematic summary of the major findings. The expression of LIFR determines the regenerative potential and trophic responsiveness of RGCs. Compared to neonatal RGCs, LIFR is reduced in adult RGCs and is further reduced in adult RGCs following optic nerve injury. A pro-regenerative treatment (*e.g.*, sterile intraocular inflammation) restores its expression to the normal mature level (*top*). LIFR overexpression in RGCs (intraocular AAV2-LIFR) promotes optic nerve regeneration independent of the ligand, LIF, and amplifies the effects of unrelated growth factors (*e.g.*, IGF1, SDF1, Ocm: *bottom left*). In contrast, LIFR knockdown abolishes regeneration induced by potent stimuli such as Pten deletion, intraocular zymosan, or miRNA-21 (*bottom center*). A truncation of extracellular domain of LIFR (LIFRΔ) also promotes regeneration (*bottom righ*t).

Finally, we characterized the expression of LIFR in sections of a postmortem human retina from a 17-year-old male. Immunohistochemistry revealed abundant LIFR expression in RBPMS-positive RGCs in the human retina (Figure 9).

**Figure 9.**
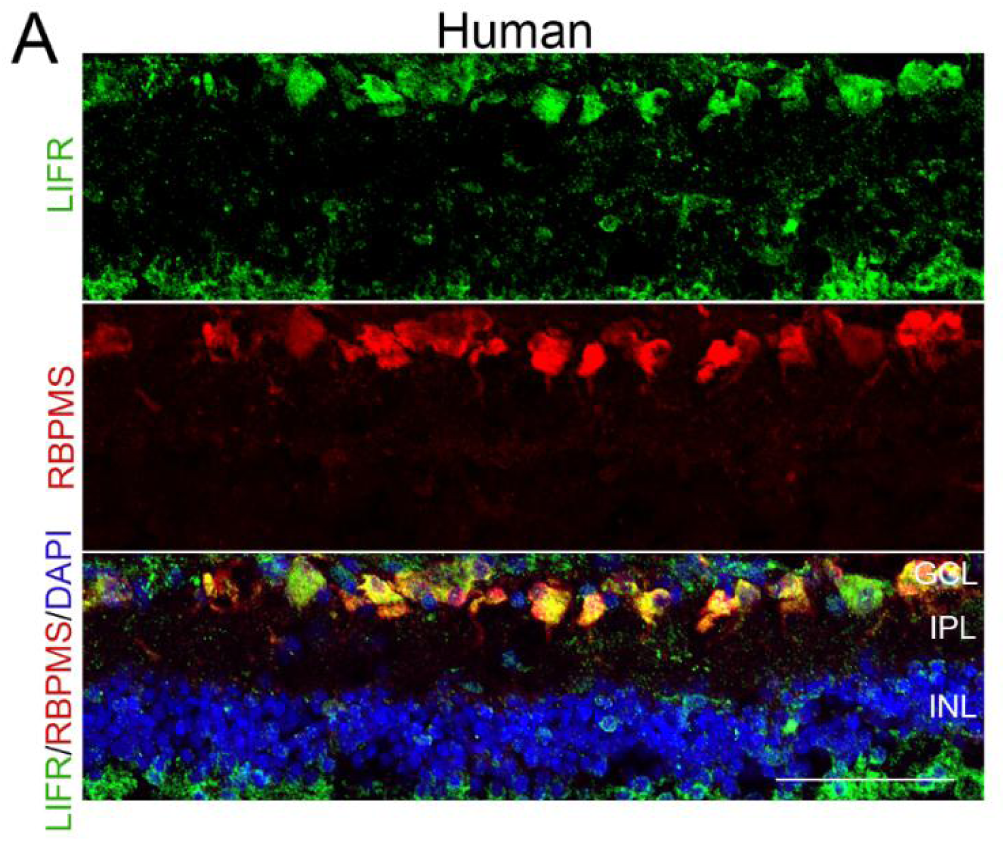
LIFR expression in human RGCs Cross-section from a human post-mortem retina immunostained for LIFR (*green*), RBPMS (*red*), and DAPI (*blue*) to visualize LIFR expression pattern in the human retina. GCL: ganglion cell layer, IPL: inner plexiform layer, INL: inner nuclear layer. Scale bar = 60 µm.

## Discussion

The present results reveal a critical role of LIFR, and particularly its intracellular domain, in the bidirectional control of optic nerve regeneration independent of the cognate ligand, LIF. Overexpression of LIFR in adult RGCs resulted in neurite outgrowth in cell culture and axon regeneration *in vivo* while strongly potentiating RGCs’ response to unrelated growth factors and to LIF itself. Conversely, LIFR deletion abrogated the effects of at least two unrelated and potent means of inducing optic nerve regeneration, Pten deletion and Zymosan-induced elevation of immune-derived trophic factors.

Mechanistically, LIFR was found to regulate intracellular signaling pathways required for axon growth, and is thus, like Pten, Socs3, and several transcription factors, a major cell-autonomous regulator of optic nerve regeneration.

LIFR is a 190 kDa protein containing an extracellular ligand-binding domain, a transmembrane domain anchoring the receptor in the cell membrane, and an intracellular domain involved in signal transduction(39). Like CNTF receptor alpha (CNTFRα)(40), LIFR functions as both a soluble and a membrane-bound receptor that forms a complex with glycoprotein 130 (gp130)(41), a receptor subunit that is shared by other cytokine receptors in the interleukin-6 (IL-6) family(42, 43). LIF binds with high affinity to the receptor complex, forming a heterodimeric complex that activates multiple signal transduction pathways(44–46). However, our RNA-seq analysis of adult RGCs revealed that LIFR regulates signal transduction pathways and gene expression in the absence of LIF. Whereas increasing LIFR expression is sufficient to increase activity of the MAP kinase pathway and expression of other growth factors, receptors, and groups of genes related to axonogenesis, growth cone formation, cytoskeletal organization, and RGCs’ response to cytokines and to myelin-associated inhibitors of axon growth independent of its cognate ligand, whereas LIFR deletion decreases the activity of both the MAPK and AKT signaling pathways, with transcriptional effects that are largely opposite to those of LIFR overexpression.

In addition to LIFR, adult RGCs express other receptors that participate in controlling axon regeneration, including ArmC10(10), CCR5 (8), CXCR4 (9), and IGF1R(13). Brain-derived neurotrophic factor (BDNF) antagonizes Zymosan-induced optic nerve regeneration(47), although virally-mediated expression of BDNF and its receptor, TrkB, enhances RGC survival(48) as reported by our lab and others. However, to our knowledge, LIFR is unique in its central role in enabling optic nerve regeneration, an effect that is independent of its cognate ligand.

During CNS development, LIFR is essential for modulating the differentiation and maturation of neuronal and non-neuronal cells(49–52) and can alleviate demyelination by enhancing oligodendrocyte survival after CNS injury(53, 54). However, whether the ligand, LIF, and the related neurokine CNTF are sufficient to induce optic nerve regeneration under normal circumstances is questionable. In agreement with several other studies, we found that, whereas CNTF by itself, has little effect on optic nerve regeneration(8, 47, 55, 56), CNTF gene therapy has robust effects (33, 46, 57–59) as a consequence of amplifying the virally-induced immune response and elevating levels of chemokine CCL5, the proximate inducer of axon regeneration(8). In addition, unlike the profound loss of regeneration found when LIFR is deleted in RGCs, deletion of the CNTF receptor CNTFRα still allows RGCs to regenerate axons, including in response to AAV2-CNTF(8). Regarding the ligand, LIF, whereas one group has reported positive effects on optic nerve regeneration(20, 60), we found little or no effect of LIF in the absence of receptor overexpression. However, because Zymosan-induced intraocular inflammation elevates levels of both LIF in neutrophils and LIFR in RGCs, it is possible that immune cell-derived LIF may supplement the effects of Ocm and SDF-1 in mediating inflammation-induced regeneration. Although blocking the effects of Ocm and SDF-1 fully abrogates Zymosan-induced regeneration(9), a non-critical but supportive role of LIF cannot be excluded. Finally, it is important to emphasize that, although prior studies have examined the role of LIF, none have examined the distinct ligand-independent role of LIFR *per se*.

In parallel to the developmental decline of RGCs’ growth capacity (61, 62), LIFR mRNA levels decrease with maturation, suggesting that LIFR may be one of the contributory factors in the decline of RGCs’ intrinsic growth potential. In addition, like Pten, socs3, c-myc and other transcription factors, our gain-and loss-of-function studies point to LIFR being a critical cell-autonomous regulator of optic nerve regeneration.

Finally, our finding that LIFR regulates adult RGCs’ responsiveness to the cognate ligand, unrelated trophic factors, and other therapies suggests that combinatorial therapies that include LIFR may maximize pro-regenerative outcomes in optic neuropathies and perhaps in other CNS neurodegenerative diseases, bringing us closer to therapeutic translation.

## Materials and Methods

### Animals and surgeries

The experiments used female and male C57BL/6J mice (Hunan SJA Laboratory), B6-G/R mice (Gempharmatech, Strain #: T006163) and vglut2-Cre mice (The Jackson Laboratory, strain #: 016963). All experiments were performed at the 2nd Xiangya Hospital with approval from the Institutional Animal Ethics Committee.

Surgeries for optic nerve crush and intravitreal injection were carried out under general anesthesia as described previously(63–65). Reagents that were injected into the vitreous chamber included zymosan (12.5µg/µl, 3µl/eye, Sigma, Cat. # Z4250), a yeast-derived glucan; recombinant mouse LIF protein (1 µg/eye, PeproTech, Cat. # 250-02); recombinant mouse IGF protein (1 µg/eye, PeproTech, Cat. # 250-19); mouse SDF1 protein (0.9 µg/eye, PeproTech, Cat. # 250-20A); mouse Ocm protein (90 ng/eye; lsbio, Cat. # LS-G57232); CPT-cAMP (50 µM, 3 µl/eye; Sigma, Cat. #C3912); LIF neutralizing antibody (400 ng/ml, 3 µl/eye, R&D, Cat. # ab449NA); CTF1 neutralizing antibody (400 ng/ml, 3 µl/eye, R&D, Cat. # AF438SP); OSM neutralizing antibody (3000 ng/ml, 3 µl/eye, R&D, Cat. # AF495NA); mouse IgG isotype control (Abcam, Cat. # ab37355); adeno-associated virus serotype 2 expressing LIFR or negative control sequence (AAV2-LIFR, AAV2-Ctrl); AAV2 expressing shRNA targeting LIFR, Pten or scramble sequence (AAV2-shLIFR, AAV2-shPten, AAV2-shSramble) and AAV2 expressing antisense sequences for miR-21 (AAV2-miR-21-sponge) or negative control (AAV2-Ctrl-sponge, GenePharma Co., Ltd); AAV2 expressing a truncated LIFR (AAV2-LIFRΔ) lacking the extracellular domain. Most agents were administered immediately after nerve injury, whereas the AAV vector was injected 2-3 weeks before NC to ensure the transfection based on experimental design. Cholera toxin subunit B (CTB, 2 µg/µl, Invitrogen, Cat. # C22843) was intraocularly injected two days before mice perfusion to trace regenerated axons.

### Human tissue collection

A postmortem human eye was collected from a donor without a known history of eye disease after removing the cornea with approval from the Ethics Committee of the 2^nd^ Xiangya Hospital. All processes in this study followed the tenets of the Declaration of Helsinki.

### Separation and identification of neutrophils in vitreous

To isolate neutrophils in the vitreous at different time points following intraocular zymosan, the vitreous was extracted into 500µl PBS and incubated with anti-Ly6G MicroBeads (Miltenyi, Cat. # 130-120-337) after washing (following the manufacturer’s protocol). Ly6G+ cells were captured by magnetic-activated cell sorting (MACS). To evaluate the purity of separated cells, cells were incubated with Gr-1 antibody conjugated with FITC (Miltenyi, Cat. # 130-102-338z) and purity was measured by flow cytometry using BD FACSCanto™ II. To characterize the morphology of isolated Ly6G+ cells, Giemsa staining was performed using Fast Giemsa Stain Kit (Solarbio, Cat. # G4640) according to the manufacturer’s protocol after cell fixation. Cells were observed and imaged under a LEICA CTR4000 microscope (Leica, Germany).

### FACS isolation of RGCs

Vglut2-tdTomato mice received an intravitreal injection of AAV2-LIFR, AAV2-Ctrl, AAV2-shLIFR, or AAV2-shScramble. Mice with different treatments were euthanized 14 days following viral vector injection to ensure the expression of encoded sequences. Retinas were carefully dissected and dissociated by trituration after incubation with papain (Sangon Biotech, Cat. # A003124-0100) for 12 mins at 37℃.

TdTomato+ Cells were isolated by fluorescent-activated cell sorting (FACS) using the BD FACSAria™ III instrument. Cells isolated from four retinas with the same treatment were pooled together. We typically collected ∼20,000 cells for each replication.

### qRT-PCR

Total RNA was extracted from RGCs, ly6G+ neutrophils, or retina using TRIzol (Invitrogen, Cat. # 15596026) following the manufacturer’s instructions. Complementary DNA (cDNA) was synthesized with RevertAid First Strand cDNA Synthesis Kit (Thermo Fisher, Cat. # K1621). QPCR was performed using Maxima SYBR Green/ROX qPCR Master Mix (Thermo Fisher, Cat. # K0221) and the following primers. LIFR-F: ATCGATGTGCAGTCCATGTATCQ, LIFR-R: ATTCCAAGTGTTTACATTGGCC, Gapdh-F: GGTTGTCTCCTGCGACTTCA, Gapdh-R: TGGTCCAGGGTTTCTTACTCC.

The relative expression level in the experimental group was initially normalized by the level of reference gene Gapdh and then by the control group level based on the experimental design.

Because neutrophils are uncommon in the vitreous without intraocular zymosan, absolute quantification in qPCR was used to measure the exact copy number of LIF in neutrophils using external standards. The external standard curves plotted to known concentrations were generated by performing qPCR on serial dilutions of a recombinant plasmid DNA encoding cDNA of LIF.

### Evaluation of Optic Nerve Regeneration and RGC survival

Animals were perfused transcardially with saline and 4% PFA (Biosharp, Cat. # BL539A). Optic nerves and retinas were dissected and post-fixed in 4% PFA for 1 hour at room temperature (RT). Post-fixed nerves were embedded into the O.C.T. compound (Sakura,Cat. # 4583) and cryostat-sectioned longitudinally at 14 µm after transferring to 30% sucrose solution at 4°C overnight. Axons labeled by Alexa Fluor™ 555 conjugated CTB (Invitrogen, Cat. # C22843) were counted in at least 3 sections per case at prespecified distances from the injury site as described(22, 63). Whole-mounted retinas were immunostained for βIII-tubulin (TUJ1, 1:500, Abcam, Cat. # ab18207) to evaluate RGC survival, as well as for CART (1:2000, Phoenix Pharmaceuticals, Cat. # H-003-62), Melanopsin (1:4000, ATS, Cat. # ATS-AB-N39), and SMI32 (1:300, BioLegend, Cat. # 801701) to visualize CART-positive on-off direction-selective RGCs (ooDSGCs), melanopsin-positive M1 to M5 intrinsically photosensitive RGCs (ipRGCs) and alpha RGCs (αRGCs) respectively, which was evaluated by counting stained cells in 8 pre-selected fields within each retina as described(63). Images were taken using Axio Imager M2+ApoTome.3 microscope (Zeiss, Germany).

### Adult dissociated retina culture

AAV2-LIFR or AAV2-Ctrl was intraocularly injected into vglut2-tdTomato mice (vglut2-Cre+/-crossed with ZsGreenflx/flx-STOPflx/flx-tdTomato) two weeks before culture. Retinas from vglut2-tdTomato mice were used to visualize RGCs in the dissociated retina culture system. The procedure for adult retinal culture has been described previously(11, 64). Briefly, retinas were dissected from adult vglut2-tdTomato mice and dissociated after incubating with papain (Sangon Biotech, Cat. # A003124-0100). Retinal cells were cultured in F12 medium supplemented with B-27 (Gibco, Cat. #s 17504044 17502048), L-glutamine (Gibco, Cat. # 25030081) and penicillin-streptomycin (Gibco, Cat. # 15140122). Additional recombinant LIF protein (PeproTech, Cat. # 250-02, 100 and 300 ng/ml) was added into the culture medium to test the effect of LIF on adult RGCs’ neurite outgrowth with or without intraocular AAV2-LIF. Neurite outgrowth and RGC survival were quantified at 3d and 4d after cell plating. TdTomato-positive RGCs survival and neurite outgrowth were quantified in 9 pre-selected fields per well in a 96-well plate. Each condition had 4-5 replications.

### Purified neonatal RGC culture

Retinas were isolated from postnatal day 1 (P1) vglut2-tdTomato mice and dissociated in the presence of papain. The immunopanning procedures used here have been described (12, 66). Briefly, dissociated retinal cells were seeded into a culture dish coated with anti-CD11b + CD11c antibody (1:500, Abcam, Cat. # ab1211) to remove glial cells. The remaining cells were transferred to another dish coated with anti-Thy1.1 antibody (1:20000, Sigma, Cat. # m7898) to select RGCs. Purified cells were then treated with trypsin and seeded into 96-well plates at a density of 7 × 10^4 cells per well. AdV-LIFR or the control vector was transfected into purified RGCs 1d following cell plating. Recombinant LIF protein (PeproTech, Cat. # 250-02, 100 ng/ml) was added 1d after viral vector transfection. After 3 days, images were taken using a Leica CTR4000 microscope (Leica, Germany). Neurite outgrowth and RGC survival were counted in 9 pre-selected fields per well. Each condition had 5 replications.

### Preparation and immunostaining of retinal cross-sections

Mice were perfused transcardially with saline followed by 4% PFA. The eyes were dissected and post-fixed in 4% PFA for 2 hours at RT. The eyeballs were embedded in O.C.T. compound and cryostat-sectioned at 14 µm after transferring into sucrose overnight at 4°C. Sections were blocked by 5% BSA followed by 0.5% Triton X-100 (DingGuo, Cat. # DH351-4). The primary antibodies used in this study included anti-RBPMS (1:400, Millipore, Cat. # ABN1376), anti-βIII-tubulin (1:500, Abcam, Cat. # ab18207), anti-LIFR (1:900, Abcam, Cat. # ab101228), and monoclonal Gr1 (1:500, LSBio, Cat. # LS-C112469). Sections were then incubated with appropriate fluorescent secondary antibodies and DAPI after washing three times.

Images were captured using an Axio Imager M2+ApoTome.3 microscope (Zeiss, Germany).

### 293T Cell line Culture and plasmid Transfection

Human 293T cells were cultured in Dulbecco’s Modified Eagle Medium (DMEM, Gibco, Cat. # c11995500bt) supplemented with 10% fetal bovine serum (FBS, Bovogen, Cat. # SFBS) at 37°C. Cells were seeded in 12-well plates at a density of 5×10^5 cells per well 24 hours before transfection. Plasmid transfections were performed using Lipofectamine 3000 (Invitrogen, Cat. #3000015) according to the manufacturer’s protocol.

### Dual-Luciferase Reporter Assay

MiR-21-binding sequence in wild-type 3’ untranslated region (UTR) of LIFR was inserted into the downstream of the luciferase reporter gene in the luciferase reporter vector. The miR-21 binding site-mutant of 3’-UTR LIFR was generated and inserted into another luciferase reporter vector. A vector containing wild-type or mutational binding sequence, together with a vector expressing miR-21 (miR-21 mimics) were transfected into 293T cells. Luciferase activity was analyzed by a dual-luciferase reporter assay system (Promega, Cat. # E2910) 48h after transfection.

## Statistical analysis

Values are shown as mean± SEM. Unpaired t-tests were used for comparisons between the two groups, whereas one-way ANOVA followed by Dunnett’s post hoc test was used if comparing three or more groups. All analyses were conducted using GraphPad Prism 9. *P<* 0.05 was considered as significant.

### RNA-seq Library Construction and sequencing

Total RNA was extracted from FACS-sorted RGCs using Trizol reagent (Invitrogen, CA, USA) according to the manufacturer’s instructions. The purity and concentration of the RNA were measured using the NanoDrop 2000 spectrophotometer (Thermo Fisher, USA). RNA integrity was assessed using the Agilent 2100 Bioanalyzer (Agilent Technologies, USA). RNA-seq library was prepared using the VAHTS Universal V5 RNA-seq Library Prep Kit (Vazyme Biotech, China) following the manufacturer’s protocol. Transcriptome sequencing and subsequent data analysis were conducted by OE Biotech Co., Ltd. (Shanghai, China).

### RNA-Seq data processing

Libraries were sequenced on a llumina Novaseq 6000 platform and 150 bp paired-end reads were generated. Raw reads of fastq format were processed using fastp(67) to remove low-quality reads and obtain clean reads for subsequent analysis. Clean reads were then aligned to the Genome Database using HISAT2(68)[59]. FPKM of each gene was calculated and the read counts of each gene were obtained by HTSeq-count(69)[60]. PCA analysis was performed using R (v3.2.0) to evaluate the biological duplication of samples. Differential expression analysis was performed using DESeq2(70). Genes with a q-value < 0.05 and a fold change (FC)>2 were identified as differentially expressed genes (DEGs). Hierarchical clustering of DEGs was conducted to illustrate expression patterns across different groups and samples. Based on the hypergeometric distribution, GO enrichment analysis(71) of DEGs was performed to screen the significantly enriched terms using R (v3.2.0).

## Acknowledgments

This work was supported by grants from the Natural Science Foundation of China (NSFC 82070967 and 81770930 to B.J., 82201189 to L.X.), China Hunan Provincial Science and Technology Department (No. 2023SK2029) and Hunan Outstanding Youth Natural Science Fund project (No. 2025JJ20080).

## Data and materials availability

All data are available in the main text or the supplementary materials. The RNA-seq data were deposited into the GEO database with accession number GSE274285.

