## Supplementary Figures & Legends for "Optic nerve regeneration requires the intracellular domain of LIFRa/CD118"

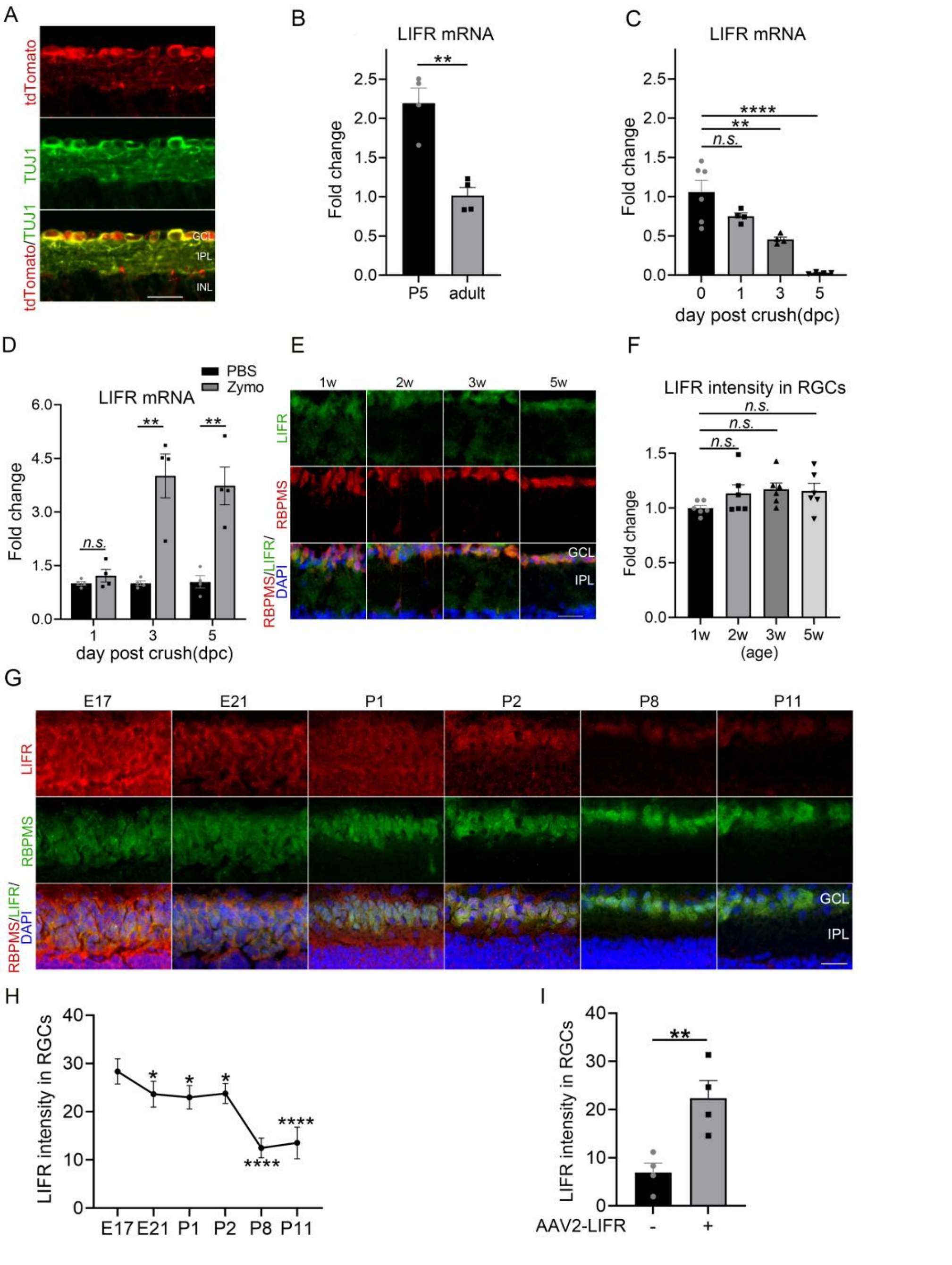


**Figure S1. LIFR intensity in RGCs during development.**

(A) Retinal cross-sections from vglut2-tdTomato mice showing genetic labeling of tdTomato (*red*) in TUJ1+ RGCs (*green*). Scale bar=60μm.

(B) LIFR mRNA expression in RGCs from postnatal day 5 (P5) and adult mice (**$p$<0.01; n = 4 samples per group).

(C) LIFR mRNA expression in RGCs from adult mice at 1, 3, and 5 days post-crush (dpc) (*n.s.* $p$>0.05, **$p$<0.01, **** $p$<0.0001; n = 4-6 samples per group).

(D) LIFR mRNA expression in adult RGCs at 1, 3, and 5 dpc with intraocular PBS or zymosan (*n.s.* $p$>0.05, **$p$<0.01; n = 4 samples per group).

(E) Retinal cross-sections from mice aged 1w, 2w, 3w and 5w immunostained for LIFR (*green*), RBPMS (*red*) and DAPI (*blue*). GCL: ganglion cell layer, IPL: inner plexiform layer. Scale bar=30 μm.

(F) Quantification of LIFR intensity in RGCs from mice aged 1w, 2w, 3w, and 5w. (*n.s.*: $p$>0.05; n=6 eyes per group).

(G) Retinal cross-sections from a rat at embryonic day 17 (E17), E21, P1, P2, P8 and P11 immunostained with LIFR (*red*), RGCs marker, RBPMS (*green*) and DAPI (*blue*). GCL: ganglion cell layer, IPL: inner plexiform layer. Scale bar = 30 μm.

(H) Quantification of LIFR intensity in RGCs from rat at E17, E21, P1, P2, P8 and P11 (*$p$<0.05, ****$p$<0.0001; n = 6 eyes per group).

(I) Quantification of LIFR intensity in adult mice two weeks following intraocular AAV2-LIFR. ( **$p$<0.01; n=4 eyes per group).

Bars show means ± SEM.

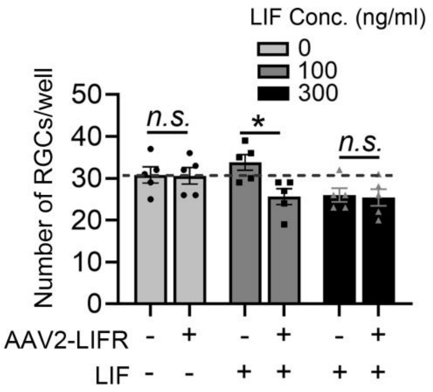


**Figure S2. LIFR overexpression does not increase adult RGC survival.**

Quantitation of RGC survival from vglut2-tdTomato mice with indicated treatments after culturing for 3 days. (*n.s.* $p>0.05$, *$p<0.05$; n = 5 wells per group).

Bars show means ± SEM.

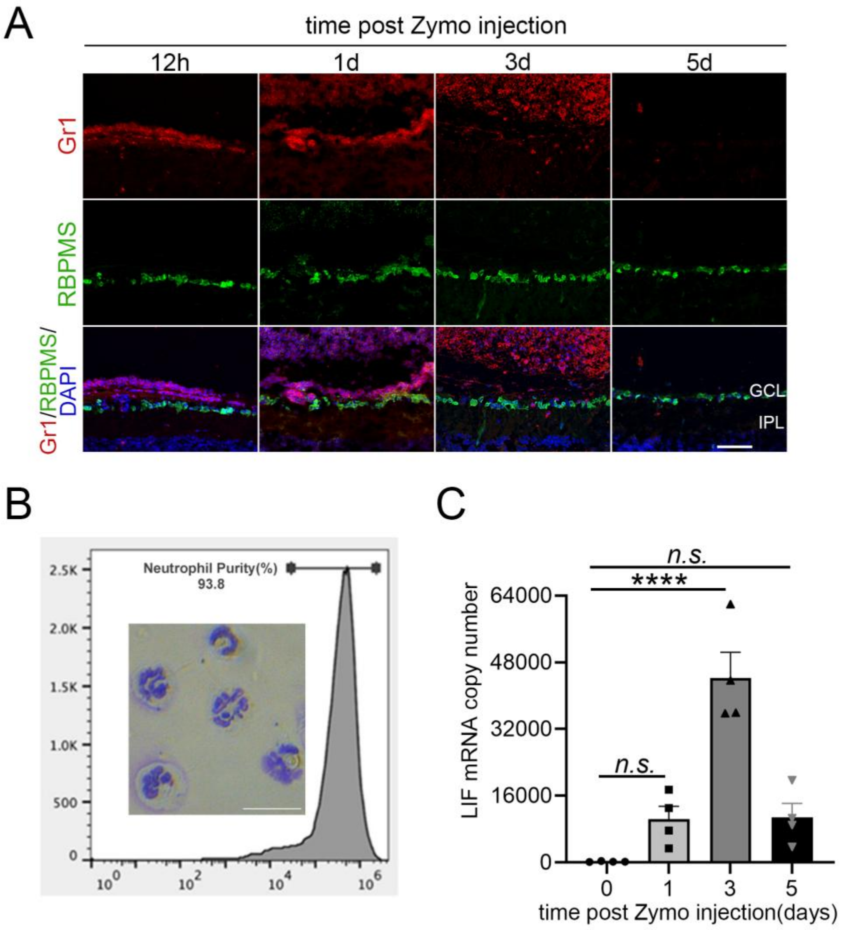


**Figure S3. Infiltrative neutrophils express LIF post intraocular zymosan**

(A) Retinal cross-sections from mice at different time points following intraocular zymosan immunostained with Gr1 (*red*), RBPMS (*green*) and DAPI (*blue*) to visualize neutrophil infiltration. GCL: ganglion cell layer, IPL: inner plexiform layer. Scale bar = 30 µm.

(B) Detection of the purification of MACS-sorted neutrophils by flow cytometry (n=3 replicates). Giemsa staining indicated the typical multilobed nuclear morphology in isolated neutrophils. Scale bar = 20 µm in inset panel.

(C) LIF mRNA level in isolated neutrophils at 1, 3, and 5d post introduction of zymosan (*n.s.* $p>0.05$, ****$p<0.0001$; n = 4 samples per group).

Bars show means ± SEM.

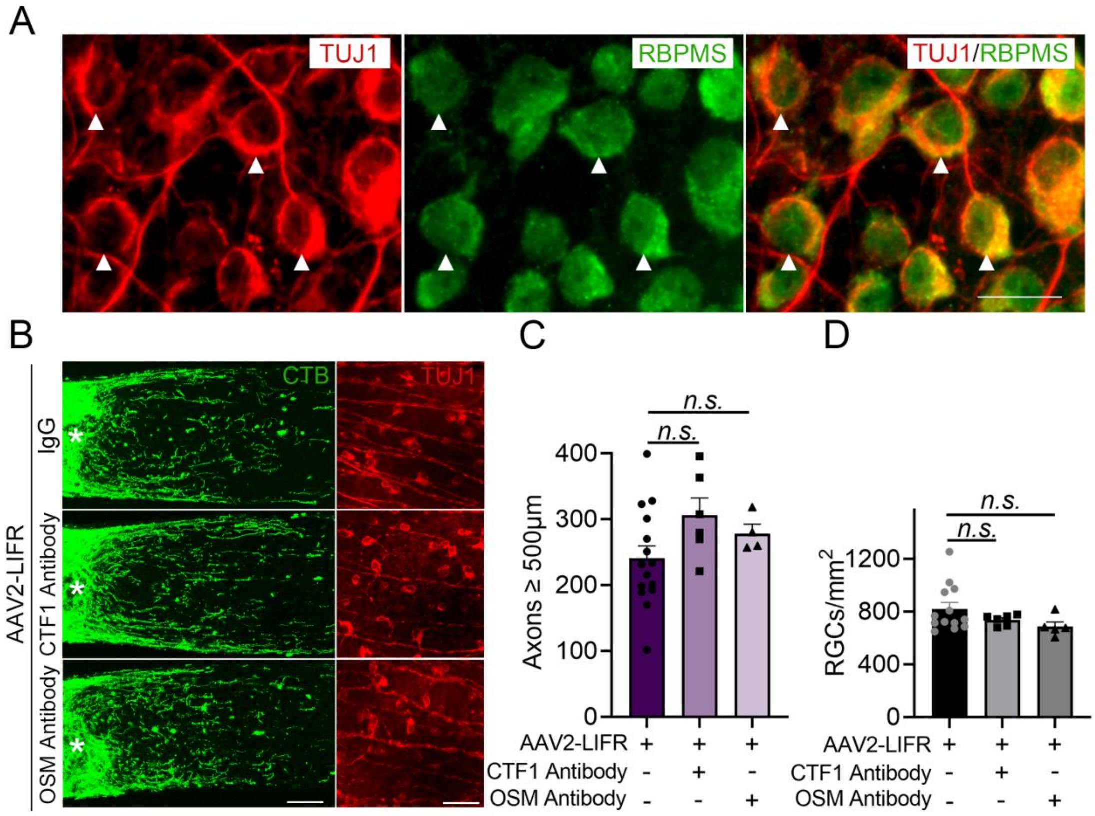


**Figure S4. The effect of LIFR overexpression is ligand-independent.**

(A) Intact whole-mounted retinas immunostained for TUJ1 (*red*) and RBPMS (*green*). TUJ1 staining was only seen in RBPMS-positive RGCs. Scale bar = 10 µm.

(B) Regenerating axons and RGC survival visualized by fluorescent CTB (*green*) and immunostaining against βIII-tubulin (*red*), respectively after intraocular AAV2-LIFR combined with CTF1 neutralizing antibody, OSM neutralizing antibody, or the isotype control. Scale bar = 100 µm in the left panel, 50 µm in the right panel.

(C) Quantification of regenerating axons (*n.s.* $p$>0.05; n = 4-15 nerves per group).

(D) Quantification of RGC survival (*n.s.* $p$>0.05; n = 5-13 eyes per group).

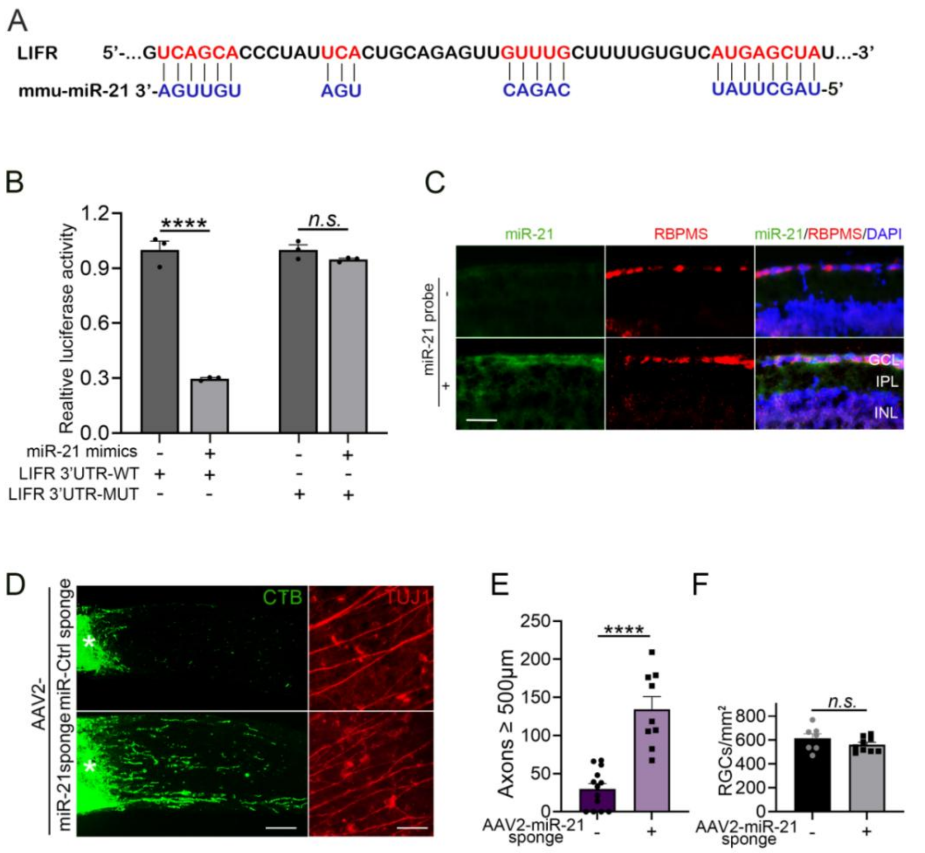


**Figure S5. AAV2-miR-21-sponge promotes axon regeneration**

(A) Colored sequences indicate the binding site between mouse miR-21 and the 3'-UTR of Lifr mRNA.

(B) Dual-luciferase reporter assay demonstrating the direct interaction between miR-21 and the 3'-UTR of LIFR mRNA. (*n.s.*: $p$>0.05, ****$p$<0.0001; n=3 wells per group).

(C) Fluorescence in situ hybridization (FISH) showing high expression of miR-21 (*green*) in RBPMS+ (*red*) RGCs.

(D) Regenerating axons and RGC survival visualized by fluorescent CTB (*green*) and immunostaining against βIII-tubulin (*red*) respectively two weeks after NC with indicated treatments. The asterisk indicates the injury site. Scale bar = 100 μm in the left panel, 50 μm in the right panel.

(E) Quantification of regenerating axons. (****$p$<0.0001; n = 9-13 nerves per group).

(F) Quantification of RGC survival. (AAV2-miR-21 sponge *vs.* AAV2-Ctrl sponge, *n.s.*: $p$>0.05; n = 7-9 eyes per group).

Bars show means ± SEM.

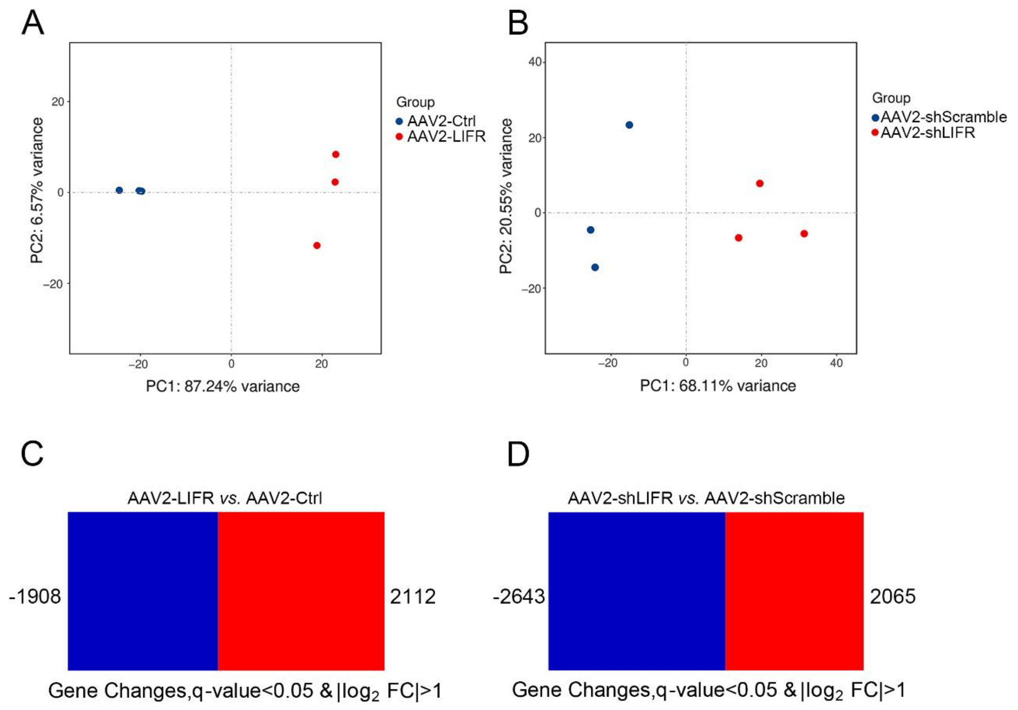


**Figure S6. Principal component analysis (PCA) and box plot**

(A, B) Principal component analysis (PCA) showing the variance in gene expression profiles in adult RGCs following transfection of AAV2-LIFR (A, AAV-LIFR *vs.* AAV-Ctrl) or AAV-shLIFR (B, AAV2-shLIFR *vs.* AAV2-shScramble).

(C, D) Box plot showing the number of DEGs ($q$<0.05, $\log_2$|FC|>1) in RGCs two weeks following intraocular AAV2-LIFR (C, AAV2-LIFR *vs.* AAV2-Ctrl) or AAV2-shLIFR (D, AAV2-shLIFR *vs.* AAV2-shScramble).

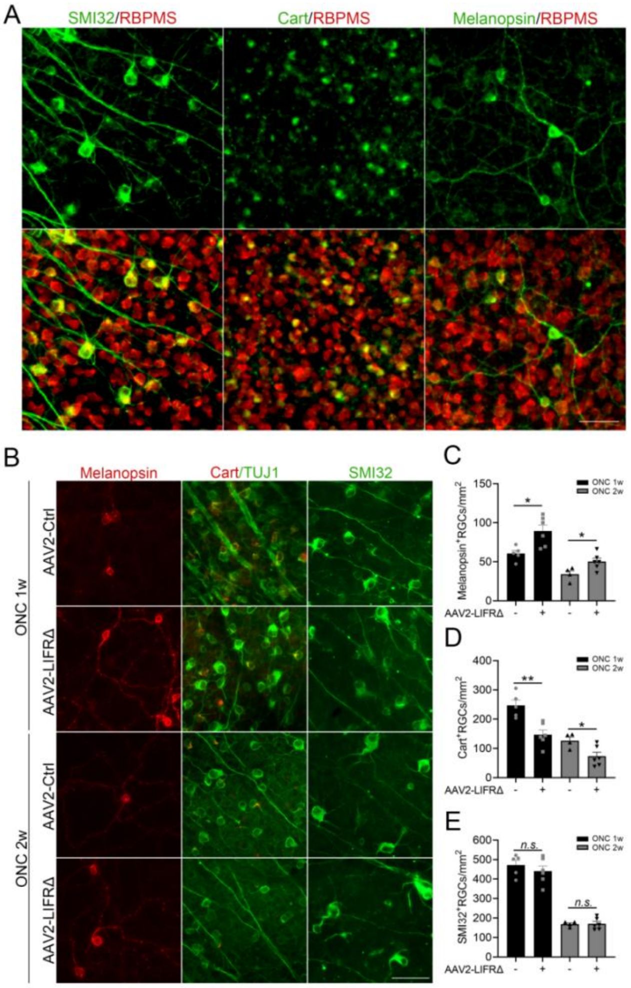
A
SMI32/RBPMS
Cart/RBPMS
Melanopsin/RBPMS
B
Melanopsin
Cart/TUJ1
SMI32
ONC 1w
ONC 2w
AAV2-Ctrl
AAV2-LIFRΔ
AAV2-Ctrl
AAV2-LIFRΔ
C
Melanopsin+RGCs/mm2
ONC 1w
ONC 2w
AAV2-LIFRΔ
D
Cart+RGCs/mm2
ONC 1w
ONC 2w
AAV2-LIFRΔ
E
SMI32+RGCs/mm2
n.s.
n.s.
ONC 1w
ONC 2w
AAV2-LIFRΔ

**Figure S7. LIFR intracellular domain alters RGC subtypes.**

(A)Whole-mounted retinas immunostained for SMI32 (*green*, left), Cart (*green*, center), or Melanopsin (*green*, right), together with the pan-RGC marker RBPMS (*red*). Scale bar = 50 µm.

(B)Whole-mounted retina stained for Melanopsin (*red*, left), Cart (*red*, middle) or SMI32 (*green*, right) to visualize ipRGCs, ooDSGCs or αRGCs, respectively at 1 and 2 weeks post-nerve injury. Scale bar = 50 µm.

(C) Quantification of ipRGCs at 1 and 2 weeks post-ONC (AAV2-LIFRΔ *vs.* AAV2-Ctrl, *$p<0.05$; n=4-6 retinas per group).

(D) Quantification of Cart+ ooDSGCs at 1 and 2 weeks post-ONC (AAV2-LIFRΔ *vs.* AAV2-Ctrl, *$p<0.05$, **$p<0.01$, n=4-6 retinas per group).

(E) Quantification of SMI32+ αRGCs at 1 and 2 weeks post-ONC (AAV2-LIFRΔ *vs.* AAV2-Ctrl, n.s.: $p>0.05$; n=4-6 retinas per group).

Bars show means ± SEM.
